# A consensus-based ensemble approach to improve transcriptome assembly

**DOI:** 10.1101/2020.06.08.139964

**Authors:** Adam Voshall, Sairam Behera, Xiangjun Li, Xiao-Hong Yu, Kushagra Kapil, Jitender S. Deogun, John Shanklin, Edgar B. Cahoon, Etsuko N. Moriyama

## Abstract

**Background:** Systems-level analyses, such as differential gene expression analysis, co-expression analysis, and metabolic pathway reconstruction, depend on the accuracy of the transcriptome. Multiple tools exist to perform transcriptome assembly from RNAseq data. However, assembling high quality transcriptomes is still not a trivial problem. This is especially the case for non-model organisms where adequate reference genomes are often not available. Different methods produce different transcriptome models and there is no easy way to determine which are more accurate. Furthermore, having alternative-splicing events exacerbates such difficult assembly problems. While benchmarking transcriptome assemblies is critical, this is also not trivial due to the general lack of true reference transcriptomes.

**Results:** In this study, we first provide a pipeline to generate a set of the benchmark transcriptome and corresponding RNAseq data. Using the simulated benchmarking datasets, we compared the performance of various transcriptome assembly approaches including both *de novo* and genome-guided methods. The results showed that the assembly performance deteriorates significantly when alternative transcripts (isoforms) exist or for genome-guided methods when the reference is not available from the same genome. To improve the transcriptome assembly performance, leveraging the overlapping predictions between different assemblies, we present a new consensus-based ensemble transcriptome assembly approach, ConSemble.

**Conclusions:** Without using a reference genome, ConSemble using four *de novo* assemblers achieved an accuracy up to twice as high as any *de novo* assemblers we compared. When a reference genome is available, ConSemble using four genome-guided assemblies removed many incorrectly assembled contigs with minimal impact on correctly assembled contigs, achieving higher precision and accuracy than individual genome-guided methods. Furthermore, ConSemble using *de novo* assemblers matched or exceeded the best performing genome-guided assemblers even when the transcriptomes included isoforms. We thus demonstrated that the ConSemble consensus strategy both for *de novo* and genome-guided assemblers can improve transcriptome assembly. The RNAseq simulation pipeline, the benchmark transcriptome datasets, and the script to perform the ConSemble assembly are all freely available from: http://bioinfolab.unl.edu/emlab/consemble/.

## Background

A high quality and comprehensive transcriptome is required in many bioinformatics workflows [1–3]. Many methods have been developed for transcriptome assembly (*e.g.*, reviewed in [4]). However, assembled transcriptomes are often incomplete, especially for non-model organisms or highly divergent strains, and further improvements are needed in assembler performance [5].

Transcriptome assembly methods can be classified into two general categories: *de novo* assemblers that generate the assembly based solely on the RNAseq data (read sets) and genome-guided assemblers that use a reference genome or transcriptome. Most of the current *de novo* assemblers, such as Trinity [6], rely on kmer decomposition of the reads, where kmers are substrings of length *k*, and de Bruijn graph construction [7]. However, with this approach, there are tradeoffs with the choice of the kmer length. For example, while shorter kmers are more likely to fully cover the transcript sequences that need to be assembled, they are also more likely to cause ambiguity in assembly graphs when repetitive sequences exist among transcripts. Due to such tradeoffs, each transcript has an optimal kmer length that facilitates the accurate reconstruction of the full-length sequence. Consequently, using different methods or even using the same method with different kmer lengths will generate different sets of contigs. Isoforms, polyploidy, multigene families, and varying levels of gene expression, all contribute to complexity in transcriptome assembly. Therefore, to obtain the complete transcriptome assembly, multiple assemblers often need to be used with a broad parameter space [8–13].

Genome-guided assemblers, such as Cufflinks [14], avoid the ambiguity in kmer assembly graphs by mapping the RNAseq data to the reference genome and clustering the reads based on genomic location [1]. However reads mapping to multiple locations within a genome can cause ambiguity [15]. When the reference genome used has any sequence divergence from the target genome, with more divergence, fewer reads can be mapped to the reference without gaps or mismatches. Each combination of read mapper and assembly algorithm handles these issues differently introducing inconsistent performance among assemblers.

While a core set of transcripts is more likely to be assembled correctly by multiple assemblers, many other transcripts may be missed depending on which specific algorithm and kmer length (for a *de novo* method) or read mapper (for a genome-guided method) are used. Through combining the results of multiple assemblers, ensemble assemblers such as EvidentialGene [16] and the method developed by Cerveau and Jackson [17] (we call their method “Concatenation”) attempt to address the limitations of individual assemblers, retaining contigs that are more likely to be correctly assembled and discarding the rest. Both of these ensemble methods filter the contigs generated by multiple assemblers (usually *de novo*) by clustering the contigs and determining the representative contig based on both the entire nucleotide and predicted protein sequences. For Concatenation, contigs that contain only portions of longer contigs (without allowing any nucleotide change) are clustered and removed. EvidentialGene clusters contigs by nucleotide similarity (98% identity by default) based on the predicted coding sequences (CDS’s) and classifies them into gene loci. From each cluster, it selects contigs including the longest CDS for the main set and those with distinct shorter CDS’s for the alternative set. Similar to Concatenation, contigs that are fragments of other contigs are removed (D. Gilbert, personal communication). Both approaches greatly reduce the number of assembled contigs by removing redundant sequences. However, there is no guarantee that the correct sequence is retained as the representative for a given cluster or that each cluster represents a unique gene. In a new genome-guided ensemble assembler, TransBorrow, after extracting reliable subpaths supported by paired-end reads from a splice graph, it merges transcripts assembled by multiple genome-guided methods and extracts reliable assembly subpaths based on the number of assemblers that detect each subpath (transcript) [18]. While the study showed superior performance of TransBorrow compared to individual genome-guided assemblers, how the performance among ensemble assemblers differs has not been examined.

To evaluate the quality of transcriptome assemblies, several methods are available, such as DETONATE [19], TransRate [15], and BUSCO [20]. The former two methods evaluate the assembly quality based on how well the assemblies are explained by the RNAseq data used. BUSCO evaluates the completeness of transcriptome assemblies by focusing on searching the universal single-copy gene sets. These methods, however, cannot measure directly the accuracy of each contig sequence. Moreover, without a benchmark transcriptome, transcripts that are not assembled (missed) cannot be quantified and the impact of false positives tends to be underestimated. In a recent study, a “real time” transcriptome was generated using the PacBio long-read sequencing technology and used as a benchmark to evaluate *de novo* assemblies generated from short-read RNAseq data obtained from the same biological samples [21]. However, obtaining good quality datasets that have been sequenced by both short- and long-read technologies, especially from many different organisms with various conditions, is not easy nor practical.

An alternative approach is to simulate the short-read sequencing based on a reference genome and generate a benchmark dataset including the reference transcriptome and the RNAseq reads. We previously used a simple simulation protocol and generated a benchmark transcriptome based on a human genome [4]. It allowed us to compare transcriptome assembly performance among different approaches. Our preliminary result showed that *de novo* transcriptome assembly can be improved by taking the consensus among multiple *de novo* assemblers.

In our present study, we generated several sets of benchmark transcriptomes with different conditions. Using these simulated benchmark datasets as well as the real RNAseq data, and using also various accuracy metrics, we performed a thorough assessment of currently available methods based on various assembly conditions. We showed that the two currently available ensemble methods that focus on thoroughness of assemblies (higher recall) retain significantly high numbers of incorrectly assembled contigs. Instead, the overlaps between different assemblies can be utilized to decrease false positives in transcriptome assembly. To demonstrate the value of a consensus approach in ensemble transcriptome assembly, we present ConSemble, a new ensemble transcriptome assembler. It combines the results from four transcriptome assemblers including both *de novo* and genome-guided methods. For *de novo* assemblers, assemblies generated using multiple kmer lengths are included. ConSemble successfully improved the accuracy of transcriptome assembly over individual tools (both *de novo* and genome-guided). Compared to other ensemble methods, ConSemble also showed consistent improvements with fewer incorrectly assembled transcripts.

## Results

### Generation of simulated benchmark RNAseq data

To understand the exact performance of different transcriptome assembly methods, we first generated simulated benchmark transcriptomes and corresponding RNAseq read sets as illustrated in the Pipeline 1 in Additional file 1. Three benchmark datasets were generated. “No0-NoAlt” and “Col0-Alt” are based on the *Arabidopsis thaliana* accessions Nossen (No-0) and Columbia (Col-0), respectively. As the simplest benchmark dataset, no alternatively spliced transcripts (isoforms) were included in “No0-NoAlt”. Another benchmark dataset was generated from the human reference genome (HG38). Table S1 in Additional file 2 shows the number and distribution of isoforms included in each dataset.

The experimental design we used to compare performance of transcriptome assembly is shown in Table S2 in Additional file 2. It allowed us to evaluate whether the transcriptome assembly performance is affected by having isoforms, and for genome-guided methods, how important the choice of the reference genome is. The assembly benchmarking process is summarized in the Pipeline 2 in Additional file 1 (see **Materials and Methods** for further details).

### Performance comparison among *de novo* transcriptome assemblers

#### Performance of individual de novo assembly methods

Using the benchmark datasets, we first compared the following four *de novo* transcriptome assemblers: Trinity [6], SOAPdenovo-Trans [22], IDBA-Tran [23], and rnaSPAdes [24, 25]. Among the three tests (Tests 1-3 in Table S2 in Additional file 2), Test 1 is the simplest where no isoforms produced by alternative splicing are included in the RNAseq dataset (No0-NoAlt). Both Tests 2 and 3 included isoforms.

All four *de novo* assemblers we tested overestimated the number of assembled contigs regardless of whether isoforms are present ranging from 119% by IDBA-Tran for Test 3 to 203% by rnaSPAdes for Test 2 (Table S3 in Additional file 2). The numbers of correctly assembled contigs were low for all four methods. Trinity produced the most accurate assembly for all datasets (*e.g.,* Precision=0.51, Recall=0.64, and *F*=0.57 for Test 1). All four assemblers performed worse when isoforms were included in the dataset (Tests 2 and 3). The isoform assembly performance was further compared in Table S4 in Additional file 2. When more than one isoform existed for a gene (Categories 3-7), often none (Category 3) or only one isoform (Category 4) could be correctly assembled by the *de novo* assemblers. Trinity was most successful among the four *de novo* assemblers in identifying multiple isoforms (higher numbers in Categories 5-7). It was the only *de novo* assembler that identified four or more isoforms (Fig. S1 right panel in Additional file 3).

Contigs assembled by these four individual assemblers are compared in Fig. 1 (left column). The 4-way intersection set included a large portion of correct contigs (40 ∼ 45%, shown in black letters) for all three datasets. It is also noteworthy that some contigs were correctly assembled only by one assembler. In contrast, while the large portion of the contigs that were assembled only by a single method were incorrect, the 4-way intersection set included only a small number of incorrect contigs (0.6∼2%, shown in red letters). These results illustrate a risk of choosing a single *de novo* assembler.

**Fig. 1.**
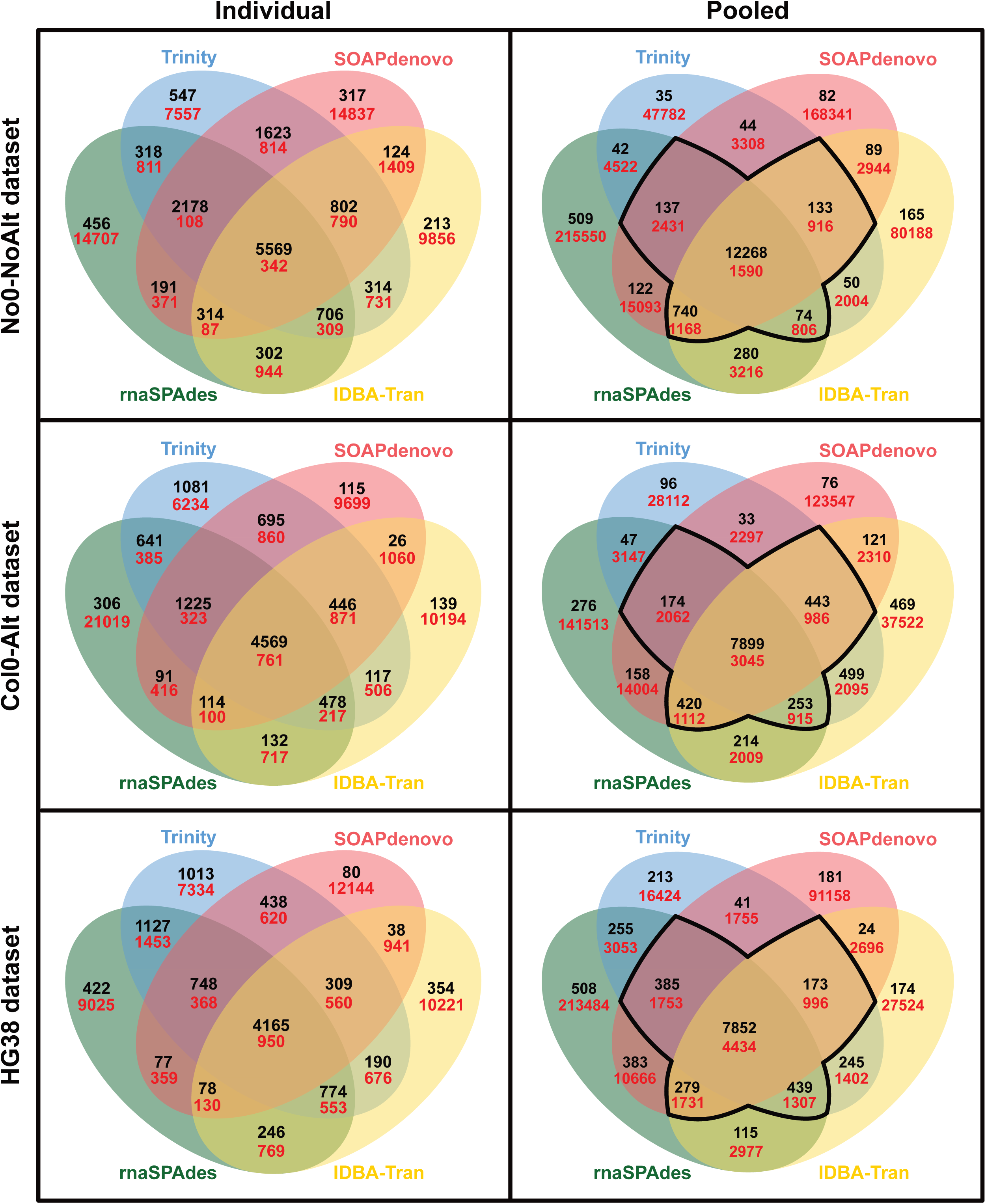
Numbers of assembled contigs shared between the four *de novo* assemblers. The numbers of correctly (black) and incorrectly (red) assembled contigs are shown. The three benchmark datasets (No0-NoAlt, Col0-Alt, and Human HG38) were assembled by the four *de novo*. Results based on individual methods using the default settings are shown under “Individual”. Results based on comparing the “pooled unique” contig sets assembled using the four methods with multiple kmer lengths are shown under “Pooled”. All contigs were compared at the protein level. The outlined region represents where the shared correct and incorrect contigs were counted for the ConSemble3+d assembly (shown as *TP* and *FP* in Table S6 in Additional file 2).

#### Development of a consensus-based de novo ensemble assembler, ConSemble

As described before, different sets of transcripts can be assembled optimally with different kmer lengths. To explore if we can increase the number of correctly assembled contigs (*TP*) with using multiple *de novo* assemblers, for each *de novo* assembler we pooled the assemblies by combining the contigs produced using multiple kmer lengths. The range and number of kmer lengths used for each method is as follows: IDBA-Tran with five kmers (20-60), SOAPdenovo-Trans with 16 kmers (15-75), rnaSPAdes with 14 kmers (19-71), and Trinity with four kmers (19-31).

As expected, this approach increased *TP* for all assemblers (Table S5 in Additional file 2; Recall>0.67 for Test 1, >0.60 for Test 2, and >0.52 for Test 3). However, it also accumulated a disproportionately large number of incorrectly assembled contigs (*FP*), greatly reducing the accuracy of the assemblies (*e.g.*, Precision=0.05 for rnaSPAdes in Test 1). When we merged all these assemblies (taking the union set of assemblies generated by all methods with multiple kmer lengths; shown as “Merged” in Table S5 in Additional file 2), the number of correctly assembled contigs increased further, although again at the cost of accumulating disproportionately more incorrectly assembled contigs. For example, for the No0-NoAlt dataset (Test 1), merging all the *de novo* assemblies produced 14,770 correctly assembled contigs (*TP*) from the 18,875 benchmark transcripts (Recall=0.78), but also had 549,859 incorrectly assembled contigs (*FP*) decreasing the Precision significantly (0.03).

When we compared these pooled contig sets assembled by the four assemblers using multiple kmers, we noted that the vast majority of the contigs that were assembled only by one or two methods were incorrect (Fig. 1, right column). However, as the number of *de novo* assemblers sharing a unique contig sequence increased, the likelihood that the contig was correctly assembled also increased. A large majority (64 ∼ 89%) of contigs produced by all four of the *de novo* assemblers were correctly assembled regardless of the test dataset, which is 70∼83% of all correctly assembled contigs. Furthermore, for all test datasets, the number of contigs that were incorrectly assembled by all four assemblers was consistently small (0.2∼1.1% of all incorrectly assembled contigs).

These observations indicate that by utilizing such consensus information, it is possible to increase the number of correctly assembled contigs and at the same time reduce the number of incorrectly assembled contigs, improving the overall assembly performance. To demonstrate this approach, we present a new consensus-based ensemble transcriptome assembly method, ConSemble, in which contigs shared by at least three of the four assemblers are retained for the final assembly (see Pipeline 3 in Additional file 1 and **Materials and Methods** for further details).

#### Performance of ConSemble compared against other de novo assembly methods

The transcriptome assembly performance of ConSemble using the four *de novo* assemblers was compared against the four individual *de novo* assemblers as well as two other ensemble methods (EvidentialGene and Concatenation) that are also based on *de novo* assemblers. Specific details for how each of these methods was run is described in **Materials and Methods**. The performance statistics and comparisons are summarized in Fig. 2 and Table S6 in Additional file 2.

**Fig. 2.**
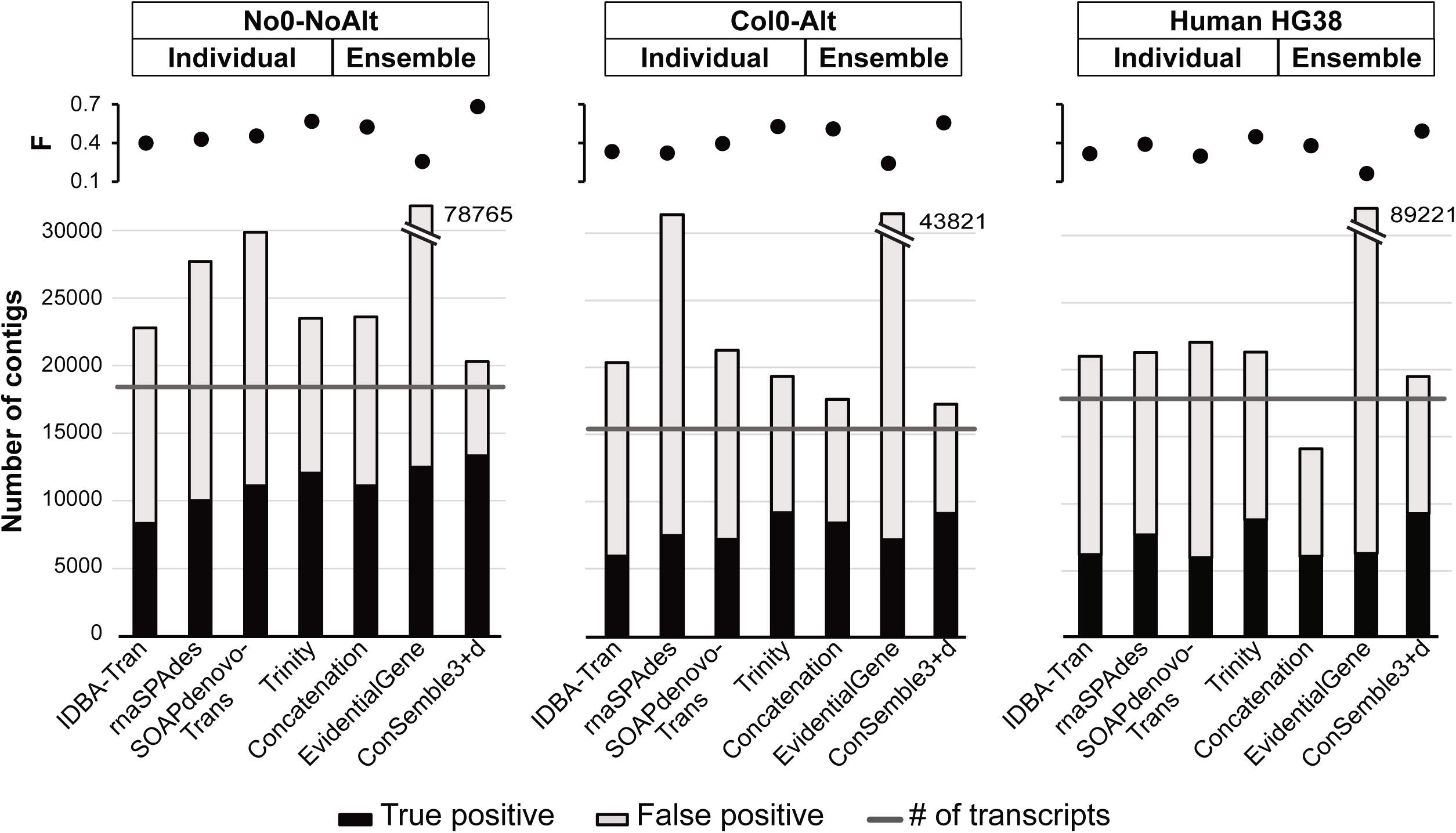
Comparison of *de novo* assembler performance on the three benchmark datasets. For the individual *de novo* assemblers, results shown were obtained with their default settings. See Tables S3 and S6 in Additional file 2 for details.

ConSemble3+d, the default assembly output of ConSemble based on four *de novo* assemblers, effectively filtered out the incorrectly assembled contigs. When isoforms were not included (the No0-NoAlt dataset), of 20,263 contigs produced by ConSemble3+d (107% of the reference), 13,352 were correctly assembled (Precision=0.66 and Recall=0.71), achieving the highest *F* score (0.68) among any *de novo* approach including both individual and ensemble methods. EvidentialGene overestimated the number of transcripts significantly (417%). While EvidentialGene recovered more benchmark transcripts (Recall=0.66) than Trinity (Recall=0.64; Table S3 in Additional file 2), the best of the individual *de novo* assemblers, many contigs were incorrectly assembled (Precision=0.16 compared to 0.51 for Trinity) leading to very low overall performance (*F*=0.26). While Concatenation produced a lower number of incorrectly assembled contigs (Precision=0.47) compared to Trinity, it did not achieve the accuracy level shown by Trinity (*F*=0.52 compared to 0.57 by Trinity).

When isoforms were included in the dataset (the Col0-Alt and HG38 datasets, Tests 2 and 3), the poor performance of the four individual *de novo* assemblers (shown in Table S3 in Additional file 2) limited the potential for these *de novo* ensemble assembly methods. ConSemble3+d slightly overestimated the number of transcripts present in the benchmark datasets (109∼112%). However, it again successfully reduced the numbers of incorrectly assembled contigs outperforming all other *de novo* methods (Precision=0.53 and 0.47, Recall=0.59 and 0.52, and *F*=0.56 and 0.49, respectively). EvidentialGene again had far more contigs than the benchmark transcriptome (283% and 335% for the Col0-Alt and HG38 datasets, respectively) resulting in very low overall performance (*F*=0.24 and 0.16, respectively). While Concatenation showed much higher Precision with lower numbers of incorrectly assembled contigs than EvidentialGene, the overall performance (*F*=0.51 and 0.38, respectively) did not exceed those obtained by using only Trinity (*F*=0.53 and 0.45, respectively). The limited ability of the most *de novo* assemblers to reconstruct multiple isoforms further limited ConSemble’s ability to accurately recover all the isoforms present in the merged assembly (Fig. S2 in Additional file 3). However, although ConSemble3+d recovered fewer isoforms especially for genes with five or more isoforms than Trinity, ConSemble3+d performed significantly better than other individual *de novo* assemblers in terms of isoform identification (Fig. S1 right panel in Additional file 3).

Among the three *de novo* ensemble assemblers, ConSemble3+d and Concatenation performs virtually the same in terms of identifying isoforms up to five (Fig. S2 in Additional file 3). For genes containing six or more isoforms, which correspond to only less than 2% of the benchmark genes, Concatenation appears to be able to identify slightly more isoforms. However, with lower Precision, Recall, and F scores with this method compared to ConSemble3+d (Table S6 in Additional file 2), it sacrifices the recovery of transcripts from genes with five or fewer isoforms.

As described before, EvidentialGene and Concatenation cluster the contigs and choose the representatives for the final assemblies. These approaches did not seem to significantly improve the overall transcriptome assembly performance compared to the best individual *de novo* assembler. However, as we observed in Table S5 in Additional file 2, the “Merged” *de novo* assembly correctly assembled up to 78% of the transcripts in the benchmark. By utilizing consensus information, the ConSemble approach successfully recovered many of these transcripts without increasing the number of incorrectly assembled contigs (*FP*), improving the overall assembly performance.

### Performance comparison among genome-guided assemblers

The ConSemble approach can also apply to genome-guided methods (Pipeline 4 in Additional file 1). Therefore, we compared the performance of genome-guided transcriptome assemblers including both individual as well as ensemble methods.

#### Performance of individual genome-guided methods

Using the benchmark datasets, we compared the following four genome-guided transcriptome assemblers: Cufflinks [14], Bayesembler [26], Scallop [27], and StringTie2 [28, 29]. The simplest experiment was to assemble the transcriptome where no alternative splicing happened (the No0-NoAlt dataset) using the same No-0 genome as a reference (Test 4 in Table S2 in Additional file 2). The genome-guided assemblers performed surprisingly poorly for this dataset (Table S7 in Additional file 2). For all assemblies, only 71∼75% of the assembled contigs were correct (shown as Precision). StringTie2 recovered the most benchmark transcripts (Recall=0.80) and Bayesembler the least (Recall=0.59).

The complexity of dealing with alternative splice forms is evident from the decreased performance of all four assemblers even when using the same genome as the reference (Tests 6 and 8 in Table S7 in Additional file 2). While Bayesembler consistently produced the fewest unique contigs, those assembled contigs were most accurate as shown by the consistently highest values of Precision (>0.54). Scallop and StringTie2, on the other hand, produced larger numbers of unique contigs recovering more benchmark transcripts as shown by the larger Recall values (>0.48).

The results were similar when STAR [30] was used as the aligner (Table S8 in Additional file 2). STAR with default parameters was shown to more accurately assign reads to the proper genomic location than Tophat2 [31]. However, we found that the assembly results based on STAR mapping were comparable to those based on Tophat2 showing no clear advantage in using one aligner over the other. Since Tophat2 remains the most widely used read mapper for genome-guided assemblies and is required for Bayesembler, only the results based on Tophat2 alignments are presented for further analyses.

The isoform assembly performance is compared further in Table S4 in Additional file 2. When more than one isoform existed for a gene (Categories 3-7), Bayesembler and Scallop recovered more isoforms (higher numbers in Categories 5-7). These two assemblers could identify all isoforms for 50% or more of the genes that have isoforms (Category 6) or for about one third of the genes that have three or more isoforms (Category 7). Among the four genome-guided assemblers, only Bayesembler identified up to seven isoforms correctly, although the success rates were not very high (Fig. S1 left panel in Additional file 3). None of the assemblers could identify all isoforms when there were more than six isoforms.

The performance of all these assemblers was considerably worse when a non-identical reference genome was used as the reference (Tests 5, 7, and 9 in Table S7 in Additional file 2). While the total numbers of contigs produced were similar to the corresponding test results with the same reference genomes (Tests 4, 6, and 8), the correctness of the contigs was greatly diminished (*e.g.*, *F*<0.34) where, on average, only 32% of the assembled contigs were correct (Precision) and only 33% of the benchmark transcripts were recovered (Recall). Of note is that all the *de novo* assemblies had higher accuracy compared to the genome-guided assemblies generated with the reference differing from the strain sequenced. Trinity, for example, assembled 41∼51% of the contigs correctly (Precision) and recovered 50∼64% of benchmark transcripts (Recall) (Table S3 in Additional file 2), while all the genome-guided assemblers showed lower than 38% for these statistics. In fact, the *de novo* assemblers performed at the level similar to those with genome-guided assemblers with the same references (Precision>0.54 and Recall>0.51; Tests 4, 6, and 8 in Table S7 in Additional file 2).

Contigs assembled by the four genome-guided assemblers are compared in Fig. 3. Similar to what we observed with the *de novo* methods, while the 4-way intersection set always included the largest numbers of correctly assembled contigs (shown in black letters), which combination of assemblers shared more correctly assembled contigs depended on the dataset and the reference genome used. Some contigs were correctly assembled only by one of the assemblers, while the majority of such contigs were false positives (incorrectly assembled, shown in red letters). Similar to what we observed with *de novo* assemblers, it illustrates the difficulty in choosing a single genome-guided assembler for any transcriptome assembly problem.

**Fig. 3.**
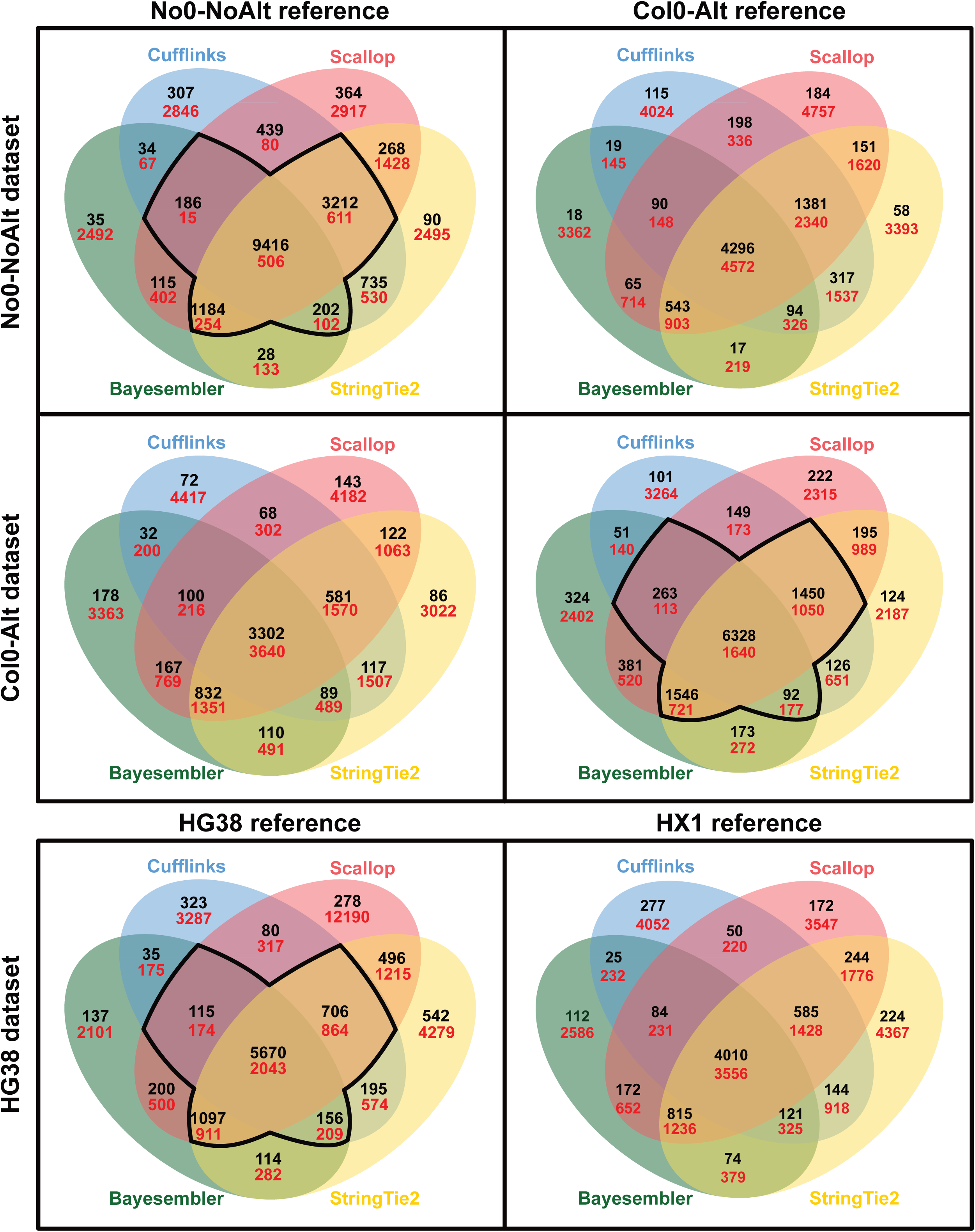
Numbers of assembled contigs shared between the four genome-guided assemblers. Rows and columns are based on the simulated RNAseq dataset and the reference genome used for the transcriptome assembly. The numbers of correctly (black) and incorrectly (red) assembled contigs are shown. All contigs were compared at the protein level. The outlined region represents where the shared correct and incorrect contigs were counted for the ConSemble3+g assembly using the same reference genomes (shown as *TP* and *FP* in Table S9 in Additional file 2).

We further compared the contigs generated by the four genome-guided assemblers with those generated by the four *de novo* assemblers. The results showed that some contigs were assembled correctly only by either genome-guided or *de novo* assemblers (Fig. 4). For example, for the No0-NoAlt dataset, 396 contigs were produced only by the *de novo* assemblers. More contigs were correctly identified uniquely by the *de novo* methods for other datasets (554 for Col0-Alt and 1,836 for Human) despite the reduced performance of the *de novo* assemblers on these datasets. It shows the advantage of using *de novo* assemblers especially when a good reference is not available.

**Fig. 4.**
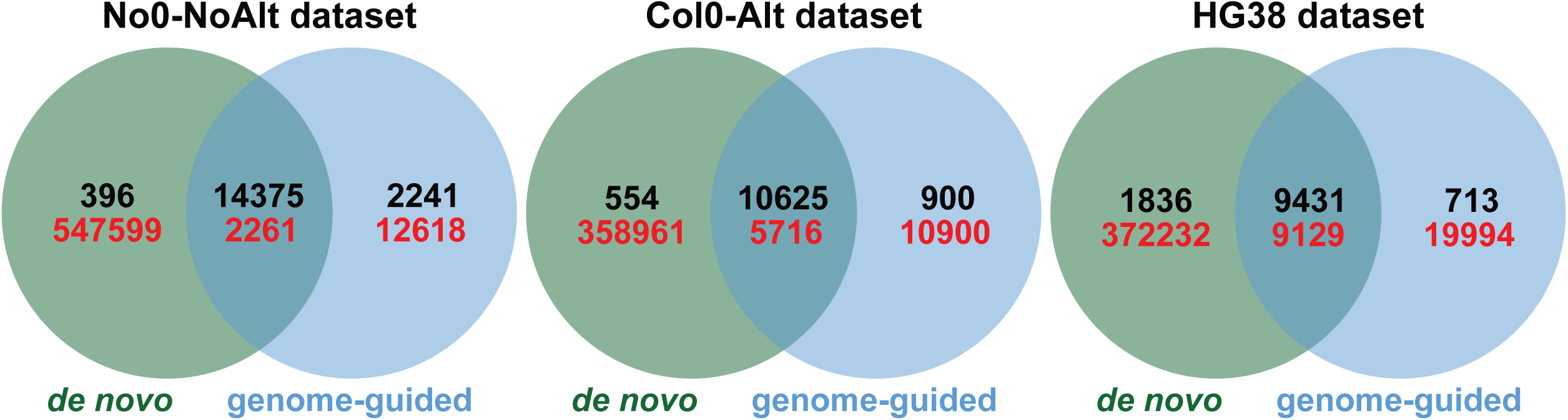
Numbers of assembled contigs shared between *de novo* and genome-guided assemblies. The “Merged” assemblies in Table S5 in Additional file 2 were used for the *de novo* assembly datasets. The genome-guided assembly is the union set of the assemblies generated by the four genome-guided methods using the same reference genomes (Tests 4, 6, and 8 in Table S2 in Additional file 2). The numbers of correctly (black) and incorrectly (red) assembled contigs are shown. All contigs were compared at the protein level.

#### Performance of ConSemble compared against other genome-guided methods

Since the individual genome-guided assemblers showed a similar trend in the overlap of correctly assembled contigs as the *de novo* methods, the consensus-based ConSemble approach with genome-guided methods is expected to improve the accuracy of the assembly. The performance of ConSemble applied to the genome-guided assemblies (ConSemble3+g) is compared with another genome-guided ensemble assembler TransBorrow [18] as well as individual genome-guided assemblers in Fig. 5 and Table S9 in Additional file 2.

**Fig. 5.**
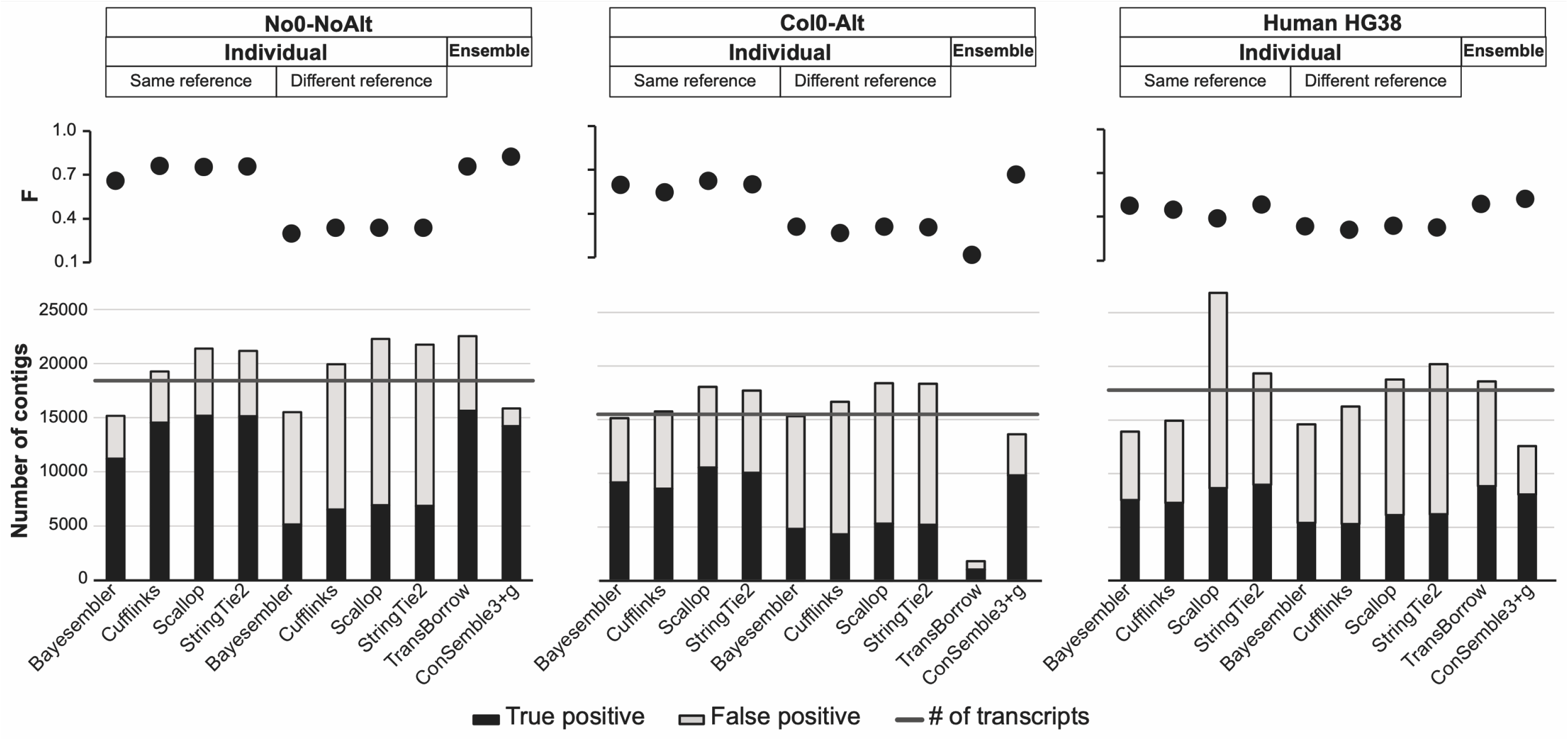
Comparison of genome-guided assembler performance on the three benchmark datasets. See Tables S7 and S9 in Additional file 2 for details.

Compared to individual genome-guided methods, TransBorrow did not show significantly different assembly performance. Moreover, TransBorrow could assemble a surprisingly small number of contigs (<12%) from the Col0-Alt dataset (Test 6). The authors mentioned about the possible issues associated to the complexity in building colored graphs depending on the assembly included [18]. The Col0-Alt dataset may have presented one such challenge. In contrast, ConSemble3+g showed significant reduction in miss-assembly (*FP*) without sacrificing correct assembly (*TP*). For the No0-NoAlt dataset, it achieved a very high Precision (0.90). For all benchmark datasets, ConSemble3+g assembly showed balanced and high accuracy (*F*=0.82, 0.67, and 0.52 for the No0-NoAlt, Col0-Alt, and HG38 datasets, respectively), which was consistently better than TransBorrow as well as individual genome-guided methods. It should be also noted that although TransBorrow recovered more correctly assembled contigs (*TP*) than ConSemble, showing higher Recall values (except for Col0-Alt), due to a larger number of incorrectly assembled contigs (*FP*), it showed lower Precision values, and hence lower overall performance (F<0.76).

For genes with large numbers of isoforms, ConSemble3+g identified more isoforms than other ensemble methods (Fig. S2 in Additional file 3). However, because many isoforms were identified in only a limited number of genome-guided assemblies (Fig. S1 left panel in Additional file 3), it also limits the ability of ConSemble3+g to identify multiple isoforms especially for genes with five or more isoforms.

### Comparison of nucleotide- and protein-level performance

In addition to the protein-level performance described above, the quality of assembled contigs at the nucleotide sequence level was examined using DETONATE [19]. DETONATE provides a reference-free model-based RSEM-EVAL score as well as reference-based scores (*F*_1_ and KC). The RSEM-EVAL metric given by DETONATE evaluates the transcriptome assembly performance by comparing the assembled nucleotide sequences to the original reads [19]. Because genome-guided assembly returns the sequence of the reference genome rather than the RNAseq data, using RSEM-EVAL on assemblies where the RNAseq data did not originate from the reference genome used can lead to poor results. Therefore, for genome-guided assemblies, only those produced using the same reference genome as the simulated RNAseq library (Tests 4, 6, and 8) were examined.

Many of the trends seen in the protein-level performance were also reflected in the nucleotide assembly quality (Table S10 in Additional file 2). In general, the assemblies with higher *F* at the protein level (Tables S3, S6, S7, and S9 in Additional file 2) also had relatively high DETONATE scores (RSEM-EVAL and KC). Among the ensemble assemblers, most often TransBorrow (except for Col0-Alt) and ConSemble3+g had the best scores in *F* and/or DETONATE scores. While EvidentialGene consistently showed the lowest *F* at the protein level as well as *F*_1_ both at the nucleotide and contig levels, it was found to perform somewhat better when the RSEM-EVAL scores were considered. As even a single nucleotide indel can have major impacts on the predicted protein sequences, it is important to evaluate performance statistics at both nucleotide and protein levels [32].

RSEM-EVAL and KC tend to provide higher scores for assemblies with longer sequences that account for, *e.g.*, more of the kmers in the RNAseq data or the reference over the precision of the contig sequence. To illustrate this issue, using the same ConSemble *de novo* assembly, we generated two other collections of contig sets. In the ConSemble3+dLong set, instead of the shortest nucleotide sequences for the default ConSemble3+d, the longest sequences were chosen as the representative contigs from those coding identical protein sequences. In the ConSemble3+dHigh set, contig sequences with the highest RSEM-EVAL scores were chosen instead. Although these three ConSemble3+d assemblies have the same protein-level *F* scores (for the same set of protein sequences), DETONATE scores were significantly different (Table S10 in Additional file 2). By choosing the shortest nucleotide sequences, ConSemble assemblies tend to truncate the untranslated regions (UTRs) of the transcripts. ConSemble3+dLong and ConSemble3+dHigh showed similar and much higher DETONATE scores indicating longer contigs are scored higher.

### Impact of using lower percent identity thresholds

The results presented so far are based on the numbers of correctly assembled contigs where the assembled sequences need to be fully 100% identical to the reference protein sequences. In practice, this is more strict than necessary for many downstream analyses. We therefore examined the impact of using lower identity thresholds on the assembly metrics.

The number of correctly assembled contigs identified at varied thresholds are shown in Fig. S3 in Additional file 3. The increase in the performance metrics was most significant when the threshold was reduced from 100% to 98%. The increase was generally proportional across all methods with the exceptions of the genome-guided assemblies using different reference genomes and EvidentialGene. With the 100% threshold, the performance of the genome-guided methods was significantly lower than *de novo* methods when different reference genomes were used. However, with less strict thresholds (<100%), their performance quickly recovered and became better than that of some of the *de novo* assemblers, although it remained lower than genome-guided assemblers using the same reference genome down to the lowest threshold (90%) used. For the No0-NoAlt dataset, EvidentialGene recovered more correctly assembled contigs with relaxed thresholds and resulted in Recall scores higher than other *de novo* methods (including both ensemble and individual methods) (Table S11 in Additional file 2). However, since EvidentialGene retained a large number of incorrectly assembled contigs (Precision<0.19 indicating only one in five contigs were correctly assembled), the accuracy level was kept very low (F<0.31). ConSemble3+d remained the best performing non-genome guided method for all three datasets. ConSemble3+g maintained the highest accuracy among all the methods we tested including individual genome-guided and *de novo* methods. These trends were consistent in the Col0-Alt and HG38 benchmark datasets as well (Tables S12 and S13 in Additional file 2).

### Assembling the real plant transcriptomes

Assembly performance was further tested using the real RNAseq data from three plant species. We included a strain of a model dicot *A. thaliana* where a high-quality genome and transcriptome information is available, another dicot *Momordica charantia* (bitter melon, a diploid), as well as the hexaploidal cotton (*Gossypium hirsutum*). This selection represents a range of ploidies, which challenges transcriptome assembly methods. In contrast to testing using simulated benchmark data, when the real RNAseq data are used, we do not know which contigs are assembled correctly or incorrectly and hence, we cannot evaluate the assembly performance using the same set of statistics. We instead used the number of genes identified from the “Eudicotyledons” dataset of Benchmarking Universal Single-Copy Orthologs (BUSCO) [20] to evaluate the thoroughness of each assembly. For *A. thaliana* Col-0, the number of contigs matching with those in the Araport11 reference transcriptome at the protein level was also counted. For the nucleotide level, the reference-free model-based RSEM-EVAL scores were included. The results are summarized in Table 1.

**Table 1.**
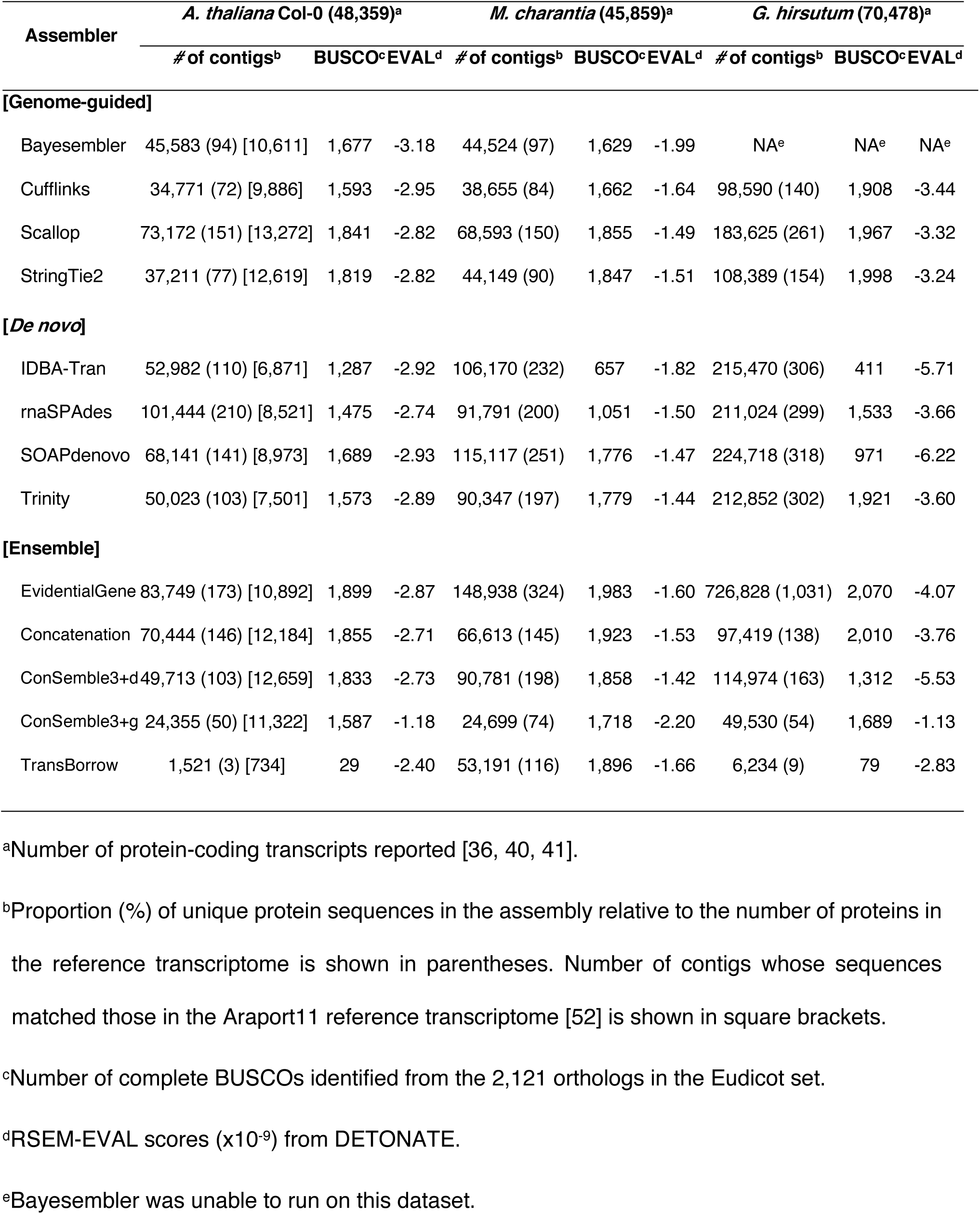
Performance analysis of real plant transcriptome assembly

For *A. thaliana* Col-0, all ensemble assemblers except TransBorrow produced more contigs that matched Araport sequences and BUSCOs and had slightly better RSEM-EVAL scores than the *de novo* assemblers. The very low performance by TransBorrow was consistent to what we observed with the Col0-Alt benchmark dataset. All ensemble assemblers had better or comparable performance than the genome-guided assemblers. While ConSemble3+g found slightly fewer BUSCOs compared to SOAPdenovo-Trans, it had the second best RSEM-EVAL following StringTie2. Among the *de novo* assembly-based ensemble methods, EvidentialGene assembled the most BUSCOs but the fewest Araport-matching contigs and had the lowest RSEM-EVAL score.

For *M. charantia*, all ensemble methods including TransBorrow recovered more BUSCOs than *de novo* assemblers. EvidentialGene, Concatenation, and TransBorrow also recovered more BUSCOs than the highest performing genome-guided method, StringTie2. Although EvidentialGene recovered the most BUSCOs, it produced a significantly more contigs (324% of the reference) indicating a high FP rate. While ConSemble3+d did not find as many BUSCOs as StringTie2 did, it had the best RSEM-EVAL score among all assemblers. Concatenation produced the fewest contigs, which was even fewer than those produced by Scallop, the best genome-guided assembler. Among the genome-guided assemblers, Scallop and StringTie2 had similar numbers of BUSCOs as well as RSEM-EVAL scores, while the number of contigs produced by StringTie2 was close to the number reported for the reference transcriptome.

For the hexaploidal cotton (*G. hirsutum*), all methods produced significantly more contigs than the 70,478 transcripts reported (except TransBorrow where the assembly largely failed). All genome-guided methods showed high numbers of BUSCOs with StringTie2 producing the most. Among the *de novo* assemblers, only Trinity produced a comparable number of BUSCOs. The remaining *de novo* assemblers produced far fewer BUSCOs despite the high numbers of contigs assembled, suggesting high false positive rates. Both EvidentialGene and Concatenation produced large numbers of BUSCOs. However, EvidentialGene produced more than ten times the number of sequences in the reference transcriptome and had a lower RSEM-EVAL score. ConSemble3+d showed a significantly smaller number of BUSCOs and one of the worst RSEM-EVAL scores. This may be due to the impact of a high ploidy [33]. It is discussed further below.

## Discussion

Despite the number of transcriptome assemblers available, there is still no single assembler or assembly strategy that performs best in all situations. Our simulation study clearly showed that every assembler correctly assembled a set of contigs while missing transcripts correctly assembled by other methods. When the reference genome was available from the same strain, the genome-guided assemblers outperformed *de novo* and most ensemble assemblers. However, if such reference sequences were not available, the performance of genome-guided assemblers dropped below that of *de novo* assemblers. Ensemble assembly approaches can overcome some of the limitations of individual assemblers. Our analysis showed that without requiring a reference genome, ensemble *de novo* methods achieved the assembly performance comparable to or higher than that of individual genome-guided methods.

Both EvidentialGene and Concatenation generally kept more of the correctly assembled contigs produced by the individual *de novo* assemblers indicated by high Recall and BUSCO scores. However, our simulation study showed that they also had many incorrectly assembled contigs that decreased the overall assembly accuracy. Instead of simply choosing representative contigs, ConSemble used consensus-based information to filter more reliable contigs. By pooling multiple assemblies, especially also pooling those generated using multiple kmer lengths for *de novo* methods, ConSemble increases the completeness of the assembled transcriptome. By taking the consensus contig set from four contig libraries, both ConSemble3+d and ConSemble3+g reduce the number of incorrectly assembled contigs significantly. This was observed in much lower numbers of incorrectly assembled contigs than those obtained by the other ensemble methods as well as all individual methods. For the three benchmark datasets, ConSemble3+d showed comparable accuracies to the genome-guided assemblers with ideal reference genomes and outperformed all individual *de novo* assemblers and the individual genome-guided assemblers without good reference genomes. Furthermore, the consensus strategy used with ConSemble allows the users to extract a set of contigs that are more likely to be correctly assembled (ConSemble4 assembly) from the rest of the contigs (ConSemble3 assembly).

Any ensemble method will necessarily be limited by the performance of the individual assemblies it is based on. If no individual method can correctly assemble a contig, no ensemble assembly can recover it. ConSemble uses the overlap of any three of the four individual assemblies (ConSemble3+). Therefore, any contigs that are correctly assembled by only one or two assemblers are omitted from the final contig set. For the three benchmark transcriptome assemblies, 10∼20% of correctly assembled contigs (1,418 of 14,770, 1,988 of 11,179, and 2,139 of 11,267 in the No0-NoAlt, Col0-Alt, and HG38 datasets, respectively) were discarded in this manner in ConSemble3+d. In Fig. S4 in Additional file 3, we examined how the assembly performance is affected by expression levels of transcripts. It shows that assembly performance is affected by the expression levels regardless of the datasets, and the *de novo* methods are affected much more significantly than the genome-guided methods. Furthermore, the ConSemble assembly (both ConSemble3+d and ConSemble3+g) performance was affected more for transcripts whose expression levels are low (*e.g.,* bin # 1-3 in Fig. S4 in Additional file 3). As we described before, especially when genes have many isoforms (*e.g.*, five or more) both ConSemble3+d and ConSemble3+g missed many of such isoforms (Fig. S2 in Additional file 3). However, as shown in “All Assemblies” in Fig. S1 in Additional file 3, many of these isoforms in fact existed in at least one of the assemblies that were used as the contig libraries for ConSemble assembly. These isoforms must have been unique to a single contig library or shared only by two contig libraries, and subsequently filtered out during the process of consensus assembly. Since the expression levels of isoforms from a gene are known to vary significantly, it is possible that the isoforms that were missed by ConSemble3+ assembly were those with low expression. This may also explain lower than expected levels of performance observed with all assemblers especially when isoforms were included in the test datasets. To recover such low expression transcripts, more refined assembly strategies beyond the current simple consensus approach need to be considered.

A similar effect is found when performance among the assemblers included in ConSemble varies widely. In the hexaploid cotton assembly, the quality of the ConSemble assembly deteriorated noticeably. This result suggests that the increased ploidy may decrease the likelihood that the true sequences are reconstructed by multiple *de novo* methods, consistent with the reduced overlap between *de novo* assemblers in simulated polyploidy benchmarks [33]. Since other ensemble approaches showed much larger BUSCOs for the cotton, it would be interesting to consider an integrated approach. Further benchmarking studies incorporating varied ploidies are necessary to determine the extent and impact of this issue.

ConSemble for *de novo* methods requires multiple different assemblers to run multiple times over a range of kmer lengths. Therefore, the availability of computational resources could become a limiting factor for deciding the number of assemblies ConSemble can utilize. However, as each assembly process is independent, these jobs can be run in parallel depending on the availability of the resources. This makes the overall assembly time roughly equivalent to the time needed for the longest individual assembly reducing the necessary computational time significantly.

## Conclusions

We presented ConSemble, a new consensus-based ensemble transcriptome assembly approach. It can be used with either *de novo* or genome-guided methods. Using ConSemble with *de novo* methods, a transcriptome assembly produced showed the quality rivals that of genome-guided assembly without requiring a reference genome and outperforms *de novo* and other ensemble strategies. ConSemble with genome-guided methods further increases the accuracy of the assembly over individual genome-guided methods. Both of *de novo* and genome-guided ConSemble assembly achieve high performance by retaining the majority of the contigs correctly assembled by individual assemblers and removing a significant number of misassembled contigs. Further work is needed for identifying more correctly assembled contigs especially when multiple lowly expressed isoforms exist or in the case of polyploidies. Improvements are needed to retain more of the contigs that are correctly assembled only by a small number of assemblers without accumulating incorrectly assembled contigs. We also showed the importance of developing realistic simulated RNAseq benchmark datasets that allow evaluating the performance of transcriptome assemblers under various conditions. It provides the only means of assessing the accuracy of the transcripts assembled directly and quantitatively.

## Materials and Methods

### Genomes and transcriptomes used

RNAseq simulation and genome-guided assemblies of the *A. thaliana* accession Nossen (No-0) were based on the assembly by Gan *et al.* [34] downloaded from the 1001 genomes project [35; accession CS6805]. RNAseq simulations and genome-guided assemblies of the *A. thaliana* accession Columbia (Col-0) were based on the TAIR reference genome (version 9) [36] and the atRTD transcriptome dataset (version 3) [37]. The atRTD dataset was chosen over the TAIR transcriptome (version 10) due to the higher prevalence of isoforms in the atRTD dataset, allowing better testing of each assembler’s ability to reconstruct alternative transcripts. The Human RNAseq simulations were based on the HG38 reference genome and transcriptome [38; GCF_000001405.39]. To evaluate the impact of using a different reference for this RNAseq library, the reads were also mapped to the HuaXia1 (HX1) reference genome (available from http://hx1.wglab.org).

Three datasets were used for the assembly of real RNAseq data. The *A. thaliana* Col-0 dataset consisting of 116M 76bp read pairs sequenced using Illumina Genome Analyzer II is from Marquez *et al.* [39], which is available at National Center for Biotechnology Information (NCBI) (SRA047499). For *M. charantia* (bitter melon), a dataset consisting of 228M 100bp read pairs was produced by sequencing of RNA samples from leaves, roots, flowers, and seeds using Illumina HiSeq 2500 (NCBI: SRR3535137, SRR3535138, SRR3535144, and SRR3535149). The reference genome was from Urasaki *et al.* [40] (NCBI: BDCS01000001–BDCS01001052). For *G. hirsutum* (upland cotton), an RNAseq dataset consisting of 117M 200bp read pairs was produced from RNA samples from leaves, roots, flowers, and seeds using Illumina HiSeq 4000 (NCBI: SRR7689126-SRR7689129). The genome assembly used as the reference is the allotetraploid L. acc. TM-1 [41; http://www.cottongen.org].

### Production of simulated benchmark transcriptomes and read sets

Simulated benchmark transcriptomes and corresponding RNAseq read sets were produced from *A. thaliana* and humans as follows. A modified Flux Simulator v1.2.1 pipeline [42] illustrated in the Pipeline 1 in Additional file 1 was used to produce ∼250M 76bp read pairs for each dataset. The No0-NoAlt transcriptome was generated based on the *A. thaliana* No-0 genomic sequence and no alternative splice events were included. The Col0-Alt transcriptome was generated based on the *A. thaliana* Col-0 genomic sequence and the version 3 atRTD transcriptome model. The expressed transcriptome in this dataset contains zero to nine alternative splice events (one to ten isoforms) per gene where only the isoforms unique at the protein level were included. The transcriptome for the Human dataset was based on the HG38 reference genome and transcriptome. It contains zero to 14 alternative splice events (1 to 15 isoforms). Only reference transcripts with full coverage of RNAseq data (all positions are required to be covered by at least one read) were included in the benchmark datasets, as transcripts without full coverage cannot be correctly assembled as a single contig. Table S1 in Additional file 2 shows the number and distribution of isoforms per transcript for each dataset. All benchmark datasets used in this study are available from: http://bioinfolab.unl.edu/emlab/consemble/

### Read processing

The read pairs generated by Flux Simulator were quality filtered using Erne-filter 2.0 [43] using the ’ultra-sensitive’ flag with a minimum average quality of q20 and in paired-end mode. The remaining reads were normalized using Khmer [44] with a kmer length of 32, an expected coverage of 50x, and in paired-end mode. As normalization works at the read level, kmer-length selection for this step has minimal impact on the unique kmers kept, and hence on the performance of the assembly.

### *De novo* assembly

*De novo* assembly was performed using Trinity 2.4.0 [6], SOAPdenovo-Trans 1.0.3 [22], IDBA-Tran 1.1.1 [23], and rnaSPAdes 3.10.0 (using the rnaspades.py script) [25]. Each of the assemblers was run using default settings as well as using multiple kmer lengths. With the default setting, IDBA-Tran assembles across multiple kmer lengths and merges the results for the final assembly. Trinity and SOAPdenovo-Trans use a single default value (25 and 23, respectively). rnaSPAdes chooses the kmer length depending on the dataset. For this study, a kmer length of 55 was chosen for both No0-NoAlt and Col0-Alt read sets and a kmer length of 37 was chosen for the HG38 set. Each method was also run with a range of kmer lengths (*k*) with increments (*i*) as follows: IDBA-Tran with *k*=20-60 and *i*=10, SOAPdenovo-Trans with *k*=15-75 and *i*=4, rnaSPAdes with *k*=19-71 and *i*=4, and Trinity with *k*=19-31 and *i*=4. The assemblers Velvet 1.2.10 [45] with Oases 0.2.08 [46] and Mira 4.0.2 [47] were excluded from this study after the preliminary analysis due to their high computational requirements (both resources and time), issues with program stability, or both.

### Genome-guided assembly

The normalized reads were first mapped to the reference genomes using Tophat2 2.0.14 with default settings [48]. Assemblies were performed using Cufflinks 2.2.1 [14], Bayesembler 1.2.0 [26], Scallop 0.10.2 [27], and StringTie2 2.0 [29]. Each assembler was executed on the default settings. Assembled contig sequences were extracted using the gtf_to_fasta module from Tophat2. The two *Arabidopsis* RNAseq datasets (No0-NoAlt and Col0-Alt) and the two human RNAseq datasets (HG38 and HX1) were assembled using each of the two reference genomes. To compare the impact of using different read mappers on assembly quality, additional assemblies were also performed using STAR 2.4.2a [30] for read mapping.

### Ensemble methods

For EvidentialGene (version 2017.03.09) [9], we chose the “okay” nucleotide contig set (okay.fa and okalt.fa) produced by the tr2aacds.pl pipeline as the final output for our comparative analysis. The assembly was performed using the same sets of four *de novo* assemblers and kmer lengths as described above resulting in 39 assemblies in total. Concatenation originally used three assemblers, Trinity, IDBA-Tran, and CLC (https://www.qiagenbioinformatics.com/), with only one kmer length each [17]. For this study, we tested all combinations of three from the four aforementioned *de novo* assemblers and kmer lengths. CLC was excluded from this study because it is only available under a commercial license. Based on the numbers of correctly and incorrectly assembled contigs, we decided to use the following three assemblies as the best combination for Concatenation: IDBA-Tran with *k*=50, Trinity with *k*=31, and rnaSPAdes with *k*=55. For TransBorrow (version 1.3) [18], the GTF files for the assemblies with the same reference genome for each of the four genome-guided assemblers (Cufflinks, Bayesembler, Scallop, and StringTie2) were manually merged into a single file. The TransBorrow assembly was performed using default parameters.

### ConSemble, a new consensus-based ensemble assembly

The Pipelines 3 and 4 in Additional file 1 summarize the process of the ConSemble transcriptome assembly approach. Four transcriptome assembly methods, either *de novo* or genome-guided, are used to generate four “contig libraries”, each containing the unique protein sequences produced by each method. For the *de novo* based assembly (Pipeline 3), each *de novo* assembler is used with a given set of multiple kmer lengths (described above), producing a total of 39 assemblies across the four methods. The assembly sets from each method are pooled to produce four contig sets. In these pooled contig sets, unique contigs are identified based on the coded protein sequences, producing four “pooled unique” contig sets. Each of the four pooled unique contig sets is used as the “contig library” from each method. For the genome-guided based assembly (Pipeline 4), the unique contig set (at the protein level) extracted from the assembled contigs using each method is used as the “contig library”. Based on the tradeoff among the performance statistics (described below), we decided to take the set of contigs that are found in the intersections of at least three contig libraries as the default output assembly. This assembly is called ConSemble3+.

The 5’ and 3’ boundaries of the transcripts are handled differently depending on the assembler. Although this does not affect the predicted open reading frame (ORF) and the coded protein sequence, it affects the overall contig length and potentially the sequence of the untranslated regions. Therefore, to minimize the impact of such differing behaviors between assemblers, we concentrated only on the longest ORFs produced by ORFfinder [49] to compare assembled contigs. When contigs with differing nucleotide sequences code entirely identical protein sequences, the one with the shortest nucleotide sequence is chosen and included in the final assembly. This is done to minimize the risk of including any chimera or over-assembled sequences. Although all redundant contigs and those with no predicted protein product are excluded from the final assembly, these contigs are saved in separate files and available for additional analyses.

ConSemble assembly can be performed based on a smaller or larger number of the assembly overlap. When only two or more assembly overlap was required (ConSemble2+d and ConSemble2+g in Tables S6 and S9 in Additional file 2), more correctly assembled contigs (*TP*) were recovered but at the cost of disproportionally more incorrectly assembled contigs (*FP*>>*TP*) leading to higher Recall but lower Precision than ConSemble3+ (*e.g.,* Recall=0.74 and 0.84 and Precision=0.27 and 0.79, respectively, for No0-NoAlt). When the 4-way consensus was used (ConSemble4d and ConSemble4g), while fewer correctly assembled contigs were recovered than ConSemble3+ (*e.g*., Recall=0.65 and 0.50 for No0-NoAlt, respectively), it reduced the number of incorrectly assembled contigs (*FP*) significantly only marginally increasing the number of missing benchmark transcripts (*FN*). This led to significantly higher Precision *(e.g*., 0.89 and 0.95 for No0-Alt, respectively) compared to ConSemble3+. Based on these results, we chose ConSemble3+ as the main output of the ConSemble pipeline. Note that, with its very high Precision, the ConSemble4 assembly can be used as part of post-processing of the ConSemble3+ assembly to obtain a contig set with the highest confidence.

ConSemble is implemented in Perl. Source codes and test data are freely available at: http://bioinfolab.unl.edu/emlab/consemble/

### Assembly benchmarking

The experimental design we used to evaluate the assembly performance is summarized in Table S2 in Additional file 2. The assembly benchmarking process is summarized in the Pipeline 2 in Additional file 1. Each contig assembly was evaluated based on the accuracy of the coded protein sequences in the ORFs predicted by ORFfinder [49]. This was done to avoid issues with variation in how the assemblers define the 3’ and 5’-ends of the contigs. Only contigs that fully covered the benchmark protein sequences without any mismatches or gaps were considered correctly assembled (true positives, TPs). Contigs with any amino acid mismatches and gaps or those corresponding to benchmark sequences only partially are considered false positives (FPs). Any transcript sequences in the benchmark dataset that were not assembled completely correctly were considered false negatives (FNs).

### Performance statistics

The overall performance of the assembly is determined as follows:

- *TP*: number of correctly assembled contigs
- *FP*: number of incorrectly assembled contigs
- *FN*: number of benchmark transcripts that are not assembled
- Recall (or sensitivity) = *TP*/(*TP*+*FN*)
- Precision = *TP*/(*TP*+*FP*)
- F-measure (*F* or *F*_1_) = 2(*TP*)/[*2(TP)+FP+FN*]

Precision shows the proportion of correctly assembled contigs relative to all assembled contigs. Recall also shows the proportion of correctly assembled contigs but relative to the number of transcripts in the reference (actual positives). F-measure is a combined metric, the harmonic mean of Precision and Recall. It therefore balances the concerns of Recall and Precision. F-measure also does not require the number of true negatives (*TN*) to be known. This is useful because a true negative is not defined for transcriptome assembly in this benchmarking. The regular accuracy score, which is defined as (*TP*+*TN*)/(*TP*+*FP*+*TN*+*FN*), also requires *TN* and cannot be calculated.

### Isoform detection performance

As described before, only isoforms that produce distinct protein sequences are considered different. This approach does not account for biologically important alternative splice events in the UTRs, which can affect protein trafficking or translation without affecting protein sequences. To simplify the analysis, the performance of the isoform detection was evaluated solely by the number of correctly assembled isoforms without considering the abundance of each isoform. Contigs that did not match completely any of the isoforms included in the benchmark dataset were counted as incorrectly assembled (FP).

### Lower identity threshold

To evaluate the effect of using % identity thresholds lower than 100% to identify TPs, identities between each assembled contig and the benchmark transcript at the translated protein level, was also calculated by dividing the edit distance by the length of the global alignment using the python library for Edlib (version 1.3.8post2) [50].

### Venn diagram generation

The contig overlap between assemblies was evaluated based on the coded protein sequences. The Venn diagrams were generated using jvenn (version 1.0) [51].

### Assembly performance at the nucleotide level

The quality of the nucleotide sequences produced by all of the assembly methods were examined using DETONATE (version 1.11) [19]. DETONATE provides a reference-free model-based RSEM-EVAL score as well as reference-based scores including the nucleotide and contig-level *F*_1_ and KC (kmer compression). RSEM-EVAL scores how well the contigs are supported by the RNAseq data as well as the compactness of an assembly based on their joint probability. The thresholds used to identify the contig-level TP to calculate *F*_1_ score (see the formula above) is 99% identity and having ≤1% of insertions and deletions. The KC score is a combination of the weighted kmer recall and inverse compression rate, the latter of which penalizes large assemblies.

### BUSCO

The thoroughness of the assemblies generated from the real plant RNAseq libraries was assessed based on the number of complete genes identified from the Eudicotyledons obd10 dataset of BUSCO (version 3.1.0) [20].

### Transcript expression level analysis

To determine the impact of expression level on transcript assembly, the benchmark transcriptome was divided into ten equally sized bins by the expression level. The proportion of transcripts in each bin correctly assembled is determined by the number of benchmark transcripts with at least one exact match (full length with no gaps or mismatches) in an assembly divided by the total number of transcripts in the bin.

## Declarations

### Ethics approval and consent to participate

Not applicable.

### Consent for publication

Not applicable.

### Availability of data and materials

The source codes, benchmark datasets, and other datasets used and analyzed in this study are available from: http://bioinfolab.unl.edu/emlab/consemble/.

### Author’s contributions

AV and ENM conceived and designed the research. AV wrote and tested the programs. XL, XHY, JS, and EBC generated RNAseq data. AV, SB, KK, and ENM analyzed and interpreted the results. JSD also interpreted the results. AV and ENM wrote and revised the manuscript. All authors read, discussed, and approved the manuscript.

## Acknowledgements

We would like to thank Dr. Don Gilbert (Indiana University) for his discussions and critical feedback regarding EvidentialGene and its performance.

## Additional files

**Additional file 1**

Format: PDF
File content: Pipelines 1-4.

**Additional file 2**

Format: PDF
File content: Supplementary Tables S1-S13.

**Additional file 3**

Format: PDF
File content: Supplementary Figs. S1-S4.

