## Additional file 1 for "A consensus-based ensemble approach to improve transcriptome assembly"

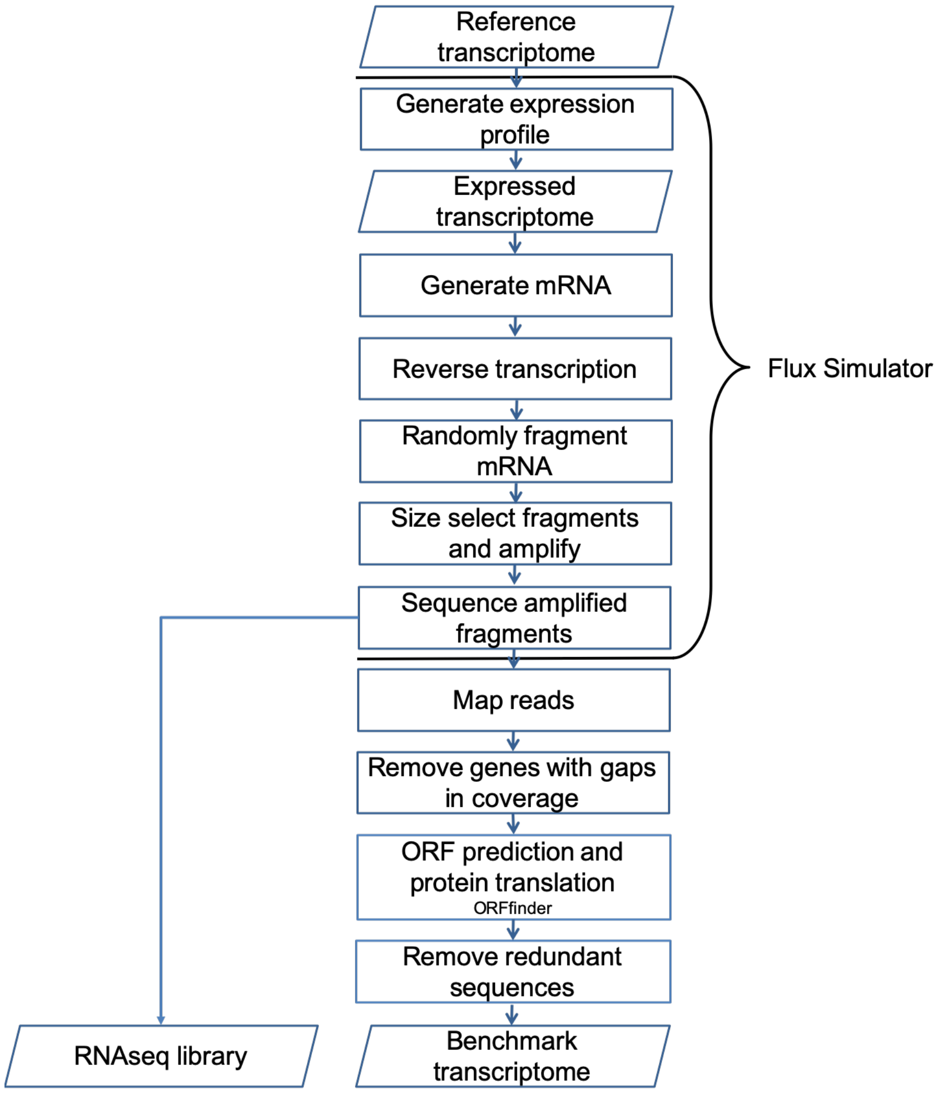


**Pipeline 1. Pipeline for generating the benchmark transcriptome and RNAseq dataset using Flux Simulator.** The expression profile is generated from the reference transcriptome and genome using the default parameters for Flux Simulator. The expressed transcripts are fragmented using a uniform random distribution, and size selected for an average length of 300bp and a standard deviation of 150bp. Fragments under 150bp are removed. Using the 76bp paired-end read sequencing error profile for Illumina HiSeq provided with Flux Simulator, read sequencing for the remaining fragments are performed. The reads produced by Flux Simulator are mapped to the expressed transcriptome using the read coordinates. Any expressed transcripts with gaps in read coverage are removed from the benchmark transcriptome. Protein-coding regions are predicted using ORFfinder and the protein sequence set corresponding to the benchmark transcriptome is produced. Only unique protein sequences were kept for the benchmark transcriptome. Note that we did not use the protein sequences available from the databases or genome projects but rather used those predicted by ORFfinder. This is to minimize the risk of using different ORF prediction and protein translation impacting the assembler benchmarking. The protein set is used for contig identification and assessment in the assembly pipeline shown in Pipeline 2. The Flux Simulator parameter file as well as sample input and output files are available at: http://bioinfolab.unl.edu/emlab/consemble

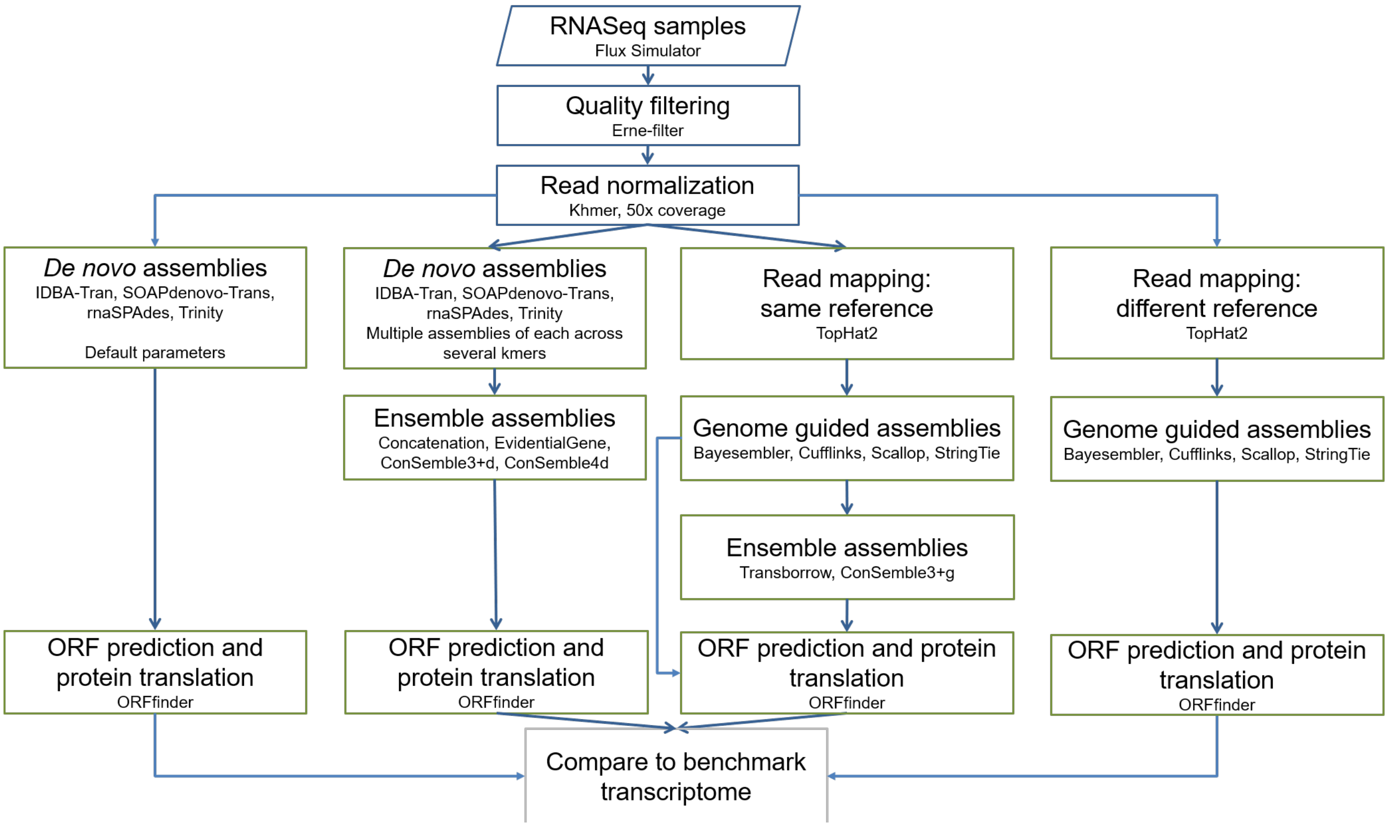


**Pipeline 2. The assembly benchmark pipeline.** See **Materials and Methods** on how each ensemble assembler was run.

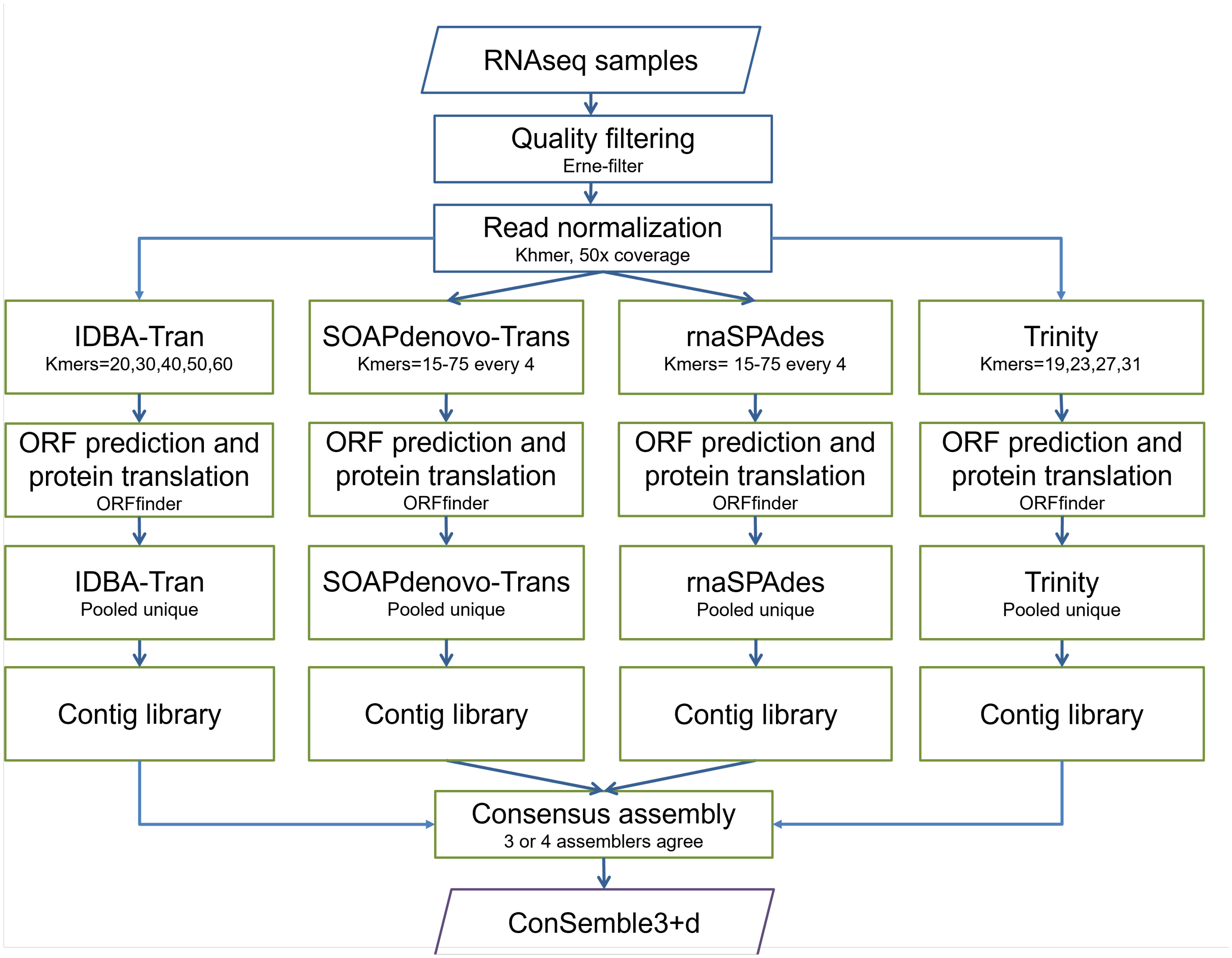


**Pipeline 3. The ConSemble pipeline for *de novo* transcriptome assembly.** After quality filtering and normalization of RNAseq data, assembly is done using each of the four *de novo* assemblers using multiple kmer values. For each assembly, protein-coding contigs are identified using ORFfinder. A unique contig set ("pooled unique") is generated for each of the *de novo* assemblers based on their protein sequences. This unique contig set forms the contig library for each of the *de novo* assembly method. Contigs that share coded protein sequences in three or more of the four contig libraries are selected for the final assembly output. This final contig set is called the "ConSemble3+d" assembly. The script for the ConSemble pipeline, sample datasets, all intermediate files, and final transcriptome outputs are available at: http://bioinfolab.unl.edu/emlab/consemble/

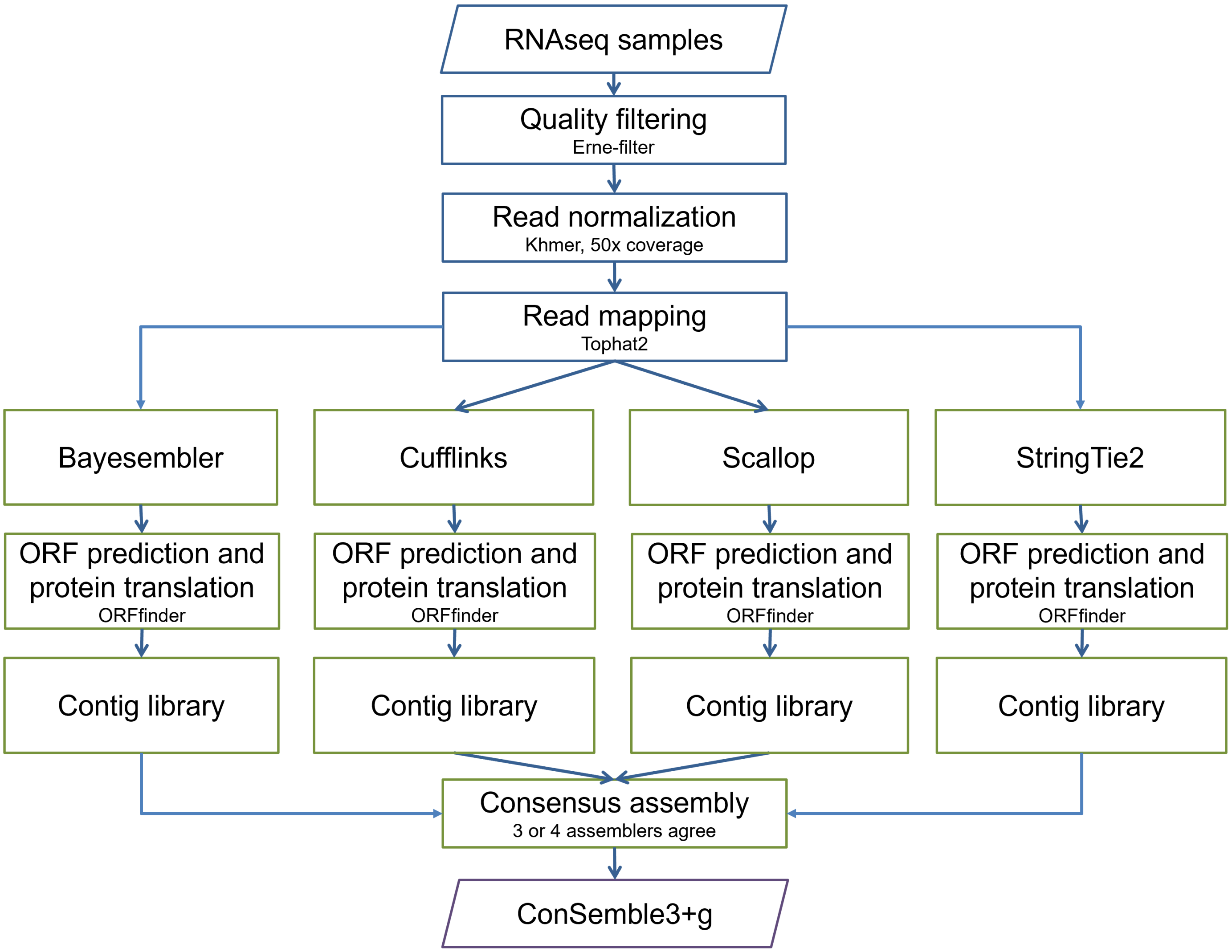


**Pipeline 4. The ConSemble pipeline for genome-guided transcriptome assembly.** After quality filtering and normalization of RNAseq data, assembly is done using each of the four genome-guided assemblers. For each assembly, protein-coding contigs are identified using ORFfinder. A unique contig set is generated for each of the *de novo* assemblers based on their protein sequences. This unique contig set forms the contig library for each method. Contigs that share coded protein sequences in at least three or more of the four contig libraries are selected for the final assembly output. This final contig set is called the "ConSemble3+g" assembly. The script for the ConSemble pipeline, sample datasets, all intermediate files, and final transcriptome outputs are available at: http://bioinfolab.unl.edu/emlab/consemble/
