## Additional file 2 for "A consensus-based ensemble approach to improve transcriptome assembly"

**Table S1. Distribution of the number of alternative transcripts in the benchmark datasets.**[a]

| # of alternative transcripts per gene[b] | # of genes | | |
|---|---|---|---|
| | **No0-NoAlt** | **Col0-Alt** | **Human HG38** |
| **1**[c] | 18,947 | 9,109 | 8481 |
| **2** | 0 | 1,915 | 2393 |
| **3** | 0 | 514 | 795 |
| **4** | 0 | 168 | 288 |
| **5** | 0 | 41 | 77 |
| **6** | 0 | 17 | 30 |
| **7** | 0 | 3 | 17 |
| **8** | 0 | 2 | 6 |
| **9** | 0 | 0 | 2 |
| **10** | 0 | 1 | 1 |
| **11** | 0 | 0 | 3 |
| **12** | 0 | 0 | 0 |
| **13** | 0 | 0 | 2 |
| **14** | 0 | 0 | 0 |
| **15** | 0 | 0 | 3 |
| **Total # of transcripts** | 18,947 | 16,071 | 18,348 |
| **Total gene length (bp)** | 30,051,975 | 24,982,520 | 28,030,410 |
| **Total # of reads**[d] | 497,448,498 | 496,589,302 | 494,422,694 |
| **Average # of reads per transcripts (minimum ~ maximum)** | 26,255 (38~8,811,100) | 26,550 (2~5,746,600) | 26,940 (2~28,287,676) |

[a]For each dataset, only the genes RNAseq reads covered the entire transcribed region are included.
[b]Only transcripts coding unique protein sequences are counted as alternative forms.
[c]These genes have no alternative splice forms.
[d]Total number of simulated RNAseq reads produced from each benchmark transcriptome.

**Table S2. Experimental design used for assessing the transcriptome assembly performance.**

| Test | Assembly type | RNAseq dataset | Reference genome | Reference type | Alternative splicing |
|---|---|---|---|---|---|
| 1 | *De novo* | No0-NoAlt | - | No | No |
| 2 | *De novo* | Col0-Alt | - | No | Yes |
| 3 | *De novo* | Human HG38 | - | No | Yes |
| 4 | Genome-guided | No0-NoAlt | No-0 | Same | No |
| 5 | Genome-guided | No0-NoAlt | Col-0 | Different | No |
| 6 | Genome-guided | Col0-Alt | Col-0 | Same | Yes |
| 7 | Genome-guided | Col0-Alt | No-0 | Different | Yes |
| 8 | Genome-guided | Human HG38 | HG38 | Same | Yes |
| 9 | Genome-guided | Human HG38 | HX1 | Different | Yes |

**Table S3. Performance analysis of *de novo* assemblers.**[a]

| Assembler | Actual[b] | Total[c] | Unique[d] | *TP* | *FP* | *FN* | Precision | Recall | *F* |
|---|---|---|---|---|---|---|---|---|---|
| Test 1: No0-NoAlt | | | | | | | | | |
| IDBA-Tran | 18,875 | 22,813 | 22,813 (120.86) | 8,344 | 14,468 | 10,531 | 0.3658 | 0.4421 | 0.4003 |
| rnaSPAdes | 18,875 | 30,478 | 27,713 (146.83) | 10,034 | 17,679 | 8,841 | 0.3621 | 0.5316 | 0.4307 |
| SOAPdenovo-Trans | 18,875 | 30,010 | 29,896 (158.28) | 11,118 | 18,758 | 7,757 | 0.3721 | 0.5890 | 0.4561 |
| Trinity | 18,875 | 23,644 | 23,519 (124.60) | 12,057 | 11,462 | 6,818 | **0.5126** | **0.6388** | **0.5688** |
| Test 2: Col0-Alt | | | | | | | | | |
| IDBA-Tran | 15,508 | 20,522 | 20,447 (131.85) | 6,021 | 14,426 | 9,487 | 0.2945 | 0.3883 | 0.3349 |
| rnaSPAdes | 15,508 | 48,241 | 31,494 (203.08) | 7,556 | 23,938 | 7,952 | 0.2399 | 0.4872 | 0.3215 |
| SOAPdenovo-Trans | 15,508 | 23,013 | 21,371 (137.81) | 7,281 | 14,090 | 8,227 | 0.3407 | 0.4695 | 0.3949 |
| Trinity | 15,508 | 20,542 | 19,409 (125.15) | 9,252 | 10,157 | 6,256 | **0.4767** | **0.5966** | **0.5299** |
| Test 3: Human HG38 | | | | | | | | | |
| IDBA-Tran | 17,669 | 20,955 | 20,954 (118.59) | 6,154 | 14,800 | 11,515 | 0.2937 | 0.3483 | 0.3187 |
| rnaSPAdes | 17,669 | 24,400 | 21,244 (120.23) | 7,637 | 13,607 | 10,032 | 0.3595 | 0.4322 | 0.3925 |
| SOAPdenovo-Trans | 17,669 | 22,499 | 22,005 (124.55) | 5,933 | 16,072 | 11,736 | 0.2697 | 0.3358 | 0.2991 |
| Trinity | 17,669 | 22,051 | 21,278 (120.43) | 8,764 | 12,514 | 8,905 | **0.4119** | **0.4960** | **0.4500** |

[a]All assemblers were run using the default settings. The best performance score for each dataset is shown in red boldface.
[b]Total number of transcripts in the benchmark transcriptome.
[c]Number of all contigs produced by the assembler.
[d]Number of unique contigs produced by the assembler. Proportion (%) of the number of transcripts in the benchmark transcriptome is shown in parentheses.

**Table S4. Comparison of the isoform assembly performance using the simulated Col0-Alt dataset.**

| Category[a] | All[b] | IDBA-Tran | SOAPdenovo-Trans | rnaSPAdes | Trinity | Bayesembler | Cufflinks | Scallop | StringTie2 |
|---|---|---|---|---|---|---|---|---|---|
| 1: No alternative splicing, assembled | 6,638 | 5,011 | 5,043 | 5,218 | 5,504 | 4,611 | 5,681 | 6,176 | 6,022 |
| 2: No alternative splicing, not assembled | 2,471 | 4,098 | 4,066 | 3,891 | 3,605 | 4,498 | 3,428 | 2,933 | 3,087 |
| 3: Alternative splicing, none assembled | 370 | 753 | 1,513 | 1,566 | 1,061 | 520 | 1,085 | 770 | 877 |
| 4: Alternative splicing, one assembled | 146 | 1,780 | 789 | 718 | 360 | 381 | 722 | 257 | 329 |
| 5: Alternative splicing, more than one assembled | 2,145 | 128 | 359 | 377 | 1,240 | 1,760 | 854 | 1,634 | 1,455 |
| 6: Alternative splicing, all assembled | 1,965 | 93 | 233 | 231 | 880 | 1,460 | 605 | 1,320 | 1,113 |
| 7: At least 3 isoforms, all assembled | 495 | 0 | 7 | 9 | 104 | 293 | 47 | 201 | 134 |

[a]Categories 1 and 2 include genes without alternative transcripts that were assembled (Category 1) and not assembled (Category 2) by each method. Categories 3-7 include those with alternative transcripts where no transcript was assembled (Category 3), only one transcript was assembled (Category 4), more than one transcript was assembled (Category 5), all transcripts were assembled (Category 6), and at least three transcripts were assembled (Category 7) by each method.

[b]Combination of all assemblies, both genome-guided and *de novo*, across all kmers. See also Fig. S1 in Additional file 3.

**Table S5. Performance analysis of *de novo* assemblers pooled across multiple kmer lengths.[a]**

| Assembler | Actual[b] | Unique[c] | *TP* | *FP* | *FN* | Precision | Recall | *F* |
|---|---|---|---|---|---|---|---|---|
| Test 1: No0-NoAlt | | | | | | | | |
| IDBA-Tran | 18,875 | 106,631 | 13,799 | 92,832 | 5,076 | 0.1294 | 0.7311 | 0.2199 |
| rnaSPAdes | 18,875 | 258,548 | 14,172 | 244,376 | 4,703 | 0.0548 | 0.7508 | 0.1022 |
| SOAPdenovo-Trans | 18,875 | 209,406 | 13,615 | 195,791 | 5,260 | 0.0650 | 0.7213 | 0.1193 |
| Trinity | 18,875 | 84,687 | 12,783 | 63,360 | 6,092 | **0.1679** | 0.6772 | **0.2691** |
| Merged[3] | 18,875 | 564,629 | 14,770 | 549,859 | 4,105 | 0.0260 | **0.7825** | 0.0506 |
| Test 2: Col0-Alt | | | | | | | | |
| IDBA-Tran | 15,508 | 60,312 | 10,318 | 49,994 | 5,190 | 0.1711 | 0.6653 | 0.2722 |
| rnaSPAdes | 15,508 | 177,248 | 9,441 | 167,807 | 6,067 | 0.0533 | 0.6088 | 0.0980 |
| SOAPdenovo-Trans | 15,508 | 158,690 | 9,324 | 149,366 | 6,184 | 0.0588 | 0.6012 | 0.1071 |
| Trinity | 15,508 | 61,208 | 9,444 | 42,660 | 6,064 | **0.1813** | 0.6088 | **0.2794** |
| Merged[c] | 15,508 | 375,854 | 11,178 | 364,676 | 4,330 | 0.0297 | **0.7208** | 0.0571 |
| Test 3: Human HG38 | | | | | | | | |
| IDBA-Tran | 17,669 | 52,368 | 9,301 | 43,067 | 8,368 | 0.1776 | 0.5264 | 0.2656 |
| rnaSPAdes | 17,669 | 249,621 | 10,216 | 239,405 | 7,453 | 0.0409 | 0.5782 | 0.0764 |
| SOAPdenovo-Trans | 17,669 | 124,507 | 9,318 | 115,189 | 8,351 | 0.0748 | 0.5274 | 0.1311 |
| Trinity | 17,669 | 40,728 | 9,603 | 31,125 | 8,066 | **0.2358** | 0.5435 | **0.3289** |
| Merged[d] | 17,669 | 392,627 | 11,267 | 381,360 | 6,402 | 0.0287 | **0.6377** | 0.0549 |

[a]The best performance score for each dataset is shown in red boldface.
[b]Total number of transcripts in the benchmark transcriptome.
[c]Number of unique contigs produced by the assembler.
[d]The Merged assembly is the union of all *de novo* assemblies across all kmer lengths

**Table S6. Performance analysis of the three ensemble transcriptome assemblers.**[a]

| Assembler | Actual[b] | Total[c] | Unique[d] | *TP* | *FP* | *FN* | Precision | Recall | *F* |
|---|---|---|---|---|---|---|---|---|---|
| Tests 1: No0-NoAlt | | | | | | | | | |
| Concatenation | 18,875 | 24,224 | 23,622 (125.15) | 11,124 | 12,498 | 7,751 | 0.4709 | 0.5894 | 0.5235 |
| EvidentialGene | 18,875 | 85,845 | 78,765 (417.30) | 12,519 | 66,246 | 6,356 | 0.1589 | 0.6633 | 0.2564 |
| ConSemble3+d | 18,875 | 20,263 | 20,263 (107.35) | 13,352 | 6,911 | 5,523 | **0.6589** | **0.7074** | **0.6823** |
| ConSemble2+d | 18,875 | 51,117 | 51,117 (275.37) | 13,979 | 37,998 | 4,896 | 0.2689 | **0.7406** | 0.3946 |
| ConSemble4d | 18,875 | 13,858 | 13,858 (73.42) | 12,268 | 1,590 | 6,607 | **0.8853** | 0.6500 | **0.7496** |
| Tests 2: Col0-Alt | | | | | | | | | |
| Concatenation | 15,508 | 18,804 | 17,717 (114,24) | 8,487 | 9,230 | 7,021 | 0.4790 | 0.5473 | 0.5109 |
| EvidentialGene | 15,508 | 46,841 | 43,821 (282.57) | 7,218 | 36,603 | 8,290 | 0.1647 | 0.4654 | 0.2433 |
| ConSemble3+d | 15,508 | 17,309 | 17,309 (111.61) | 9,189 | 8,120 | 6,319 | **0.5309** | **0.5925** | **0.5600** |
| ConSemble2+d | 15,508 | 44,243 | 44,243 (285.29) | 10,261 | 33,982 | 5,247 | 0.2319 | **0.6617** | 0.3435 |
| ConSemble4d | 15,508 | 10,944 | 10,944 (70.57) | 7,899 | 3,045 | 7,609 | **0.7218** | 0.5094 | **0.5972** |
| Tests 3: Human HG38 | | | | | | | | | |
| Concatenation | 17,669 | 15,180 | 14,061 (79.58) | 6,032 | 8,029 | 11,637 | 0.4290 | 0.3414 | 0.3802 |
| EvidentialGene | 17,669 | 65,587 | 59,221 (335.17) | 6,248 | 52,973 | 11,421 | 0.1055 | 0.3536 | 0.1625 |
| ConSemble3+d | 17,669 | 19,349 | 19,349 (109.51) | 9,128 | 10,221 | 8,541 | **0.4718** | **0.5166** | **0.4932** |
| ConSemble2+d | 17,669 | 42,961 | 42,961 (243.14) | 10,191 | 32,770 | 7,478 | 0.2372 | **0.5768** | 0.3362 |
| ConSemble4d | 17,669 | 12,286 | 12,286 (69.53) | 7,852 | 4,434 | 9,817 | **0.6391** | 0.4444 | **0.5243** |

[a]The best performance score for each dataset among Concatenation, EvidentialGene, and ConSemble3+d is shown in red boldface. When ConSemble2+d or ConSemble4d showed better performance than ConSemble3+d, such scores are shown in black boldface.

[b]Total number of transcripts in the benchmark transcriptome

[c]Number of all contigs produced by the assembler.

[d]Number of unique contigs produced by the assembler. Proportion (%) of the number of transcripts in the benchmark transcriptome is shown in parentheses.

**Table S7. Performance analysis of genome-guided assemblers using the Tophat2 aligner.**[a]

| Assembler | Actual[b] | Total[c] | Unique[d] | *TP* | *FP* | *FN* | Precision | Recall | *F* |
|---|---|---|---|---|---|---|---|---|---|
| **[No0-NoAlt]** | | | | | | | | | |
| Test 4 (Reference: No-0) | | | | | | | | | |
| Bayesembler | 18,875 | 15,297 | 15,171 (80.38) | 11,200 | 3,971 | 7,675 | 0.7383 | 0.5934 | 0.6579 |
| Cufflinks | 18,875 | 19,398 | 19,288 (102.19) | 14,531 | 4,757 | 4,344 | **0.7534** | 0.7699 | **0.7615** |
| Scallop | 18,875 | 21,853 | 21,397 (113.36) | 15,184 | 6,213 | 3,691 | 0.7096 | **0.8045** | 0.7541 |
| StringTie2 | 18,875 | 21,651 | 21,194 (112.29) | 15,135 | 6,059 | 3,740 | 0.7141 | 0.8019 | 0.7554 |
| Test 5 (Reference: Col-0) | | | | | | | | | |
| Bayesembler | 18,875 | 15,670 | 15,531 (82.28) | 5,142 | 10,389 | 13,733 | **0.3311** | 0.2724 | 0.2989 |
| Cufflinks | 18,875 | 20,133 | 19,938 (105.63) | 6,510 | 13,428 | 12,365 | 0.3265 | 0.3449 | 0.3355 |
| Scallop | 18,875 | 22,804 | 22,298 (118.14) | 6,908 | 15,390 | 11,967 | 0.3098 | 0.3660 | 0.3356 |
| StringTie2 | 18,875 | 22,275 | 21,767 (115.32) | 6,857 | 14,910 | 12,018 | 0.3150 | **0.3633** | **0.3374** |
| **[Col0-Alt]** | | | | | | | | | |
| Test 6 (Reference: Col-0) | | | | | | | | | |
| Bayesembler | 15,508 | 16,271 | 15,143 (97.65) | 9,158 | 5,985 | 6,350 | **0.6048** | 0.5905 | 0.5976 |
| Cufflinks | 15,508 | 16,232 | 15,768 (101.68) | 8,560 | 7,208 | 6,948 | 0.5429 | 0.5520 | 0.5474 |
| Scallop | 15,508 | 19,454 | 18,055 (116.42) | 10,534 | 7,521 | 4,974 | 0.5834 | **0.6793** | **0.6277** |
| StringTie2 | 15,508 | 18,897 | 17,721 (114.27) | 10,034 | 7,687 | 5,474 | 0.5662 | 0.6470 | 0.6039 |
| Test 7 (Reference: No-0) | | | | | | | | | |
| Bayesembler | 15,508 | 16,384 | 15,329 (98.85) | 4,810 | 10,519 | 10,698 | **0.3138** | 0.3102 | 0.3120 |
| Cufflinks | 15,508 | 17,303 | 16,662 (107.44) | 4,321 | 12,341 | 11,187 | 0.2593 | 0.2786 | 0.2686 |
| Scallop | 15,508 | 19,539 | 18,408 (118.70) | 5,315 | 13,093 | 10,193 | 0.2887 | **0.3427** | **0.3134** |
| StringTie2 | 15,508 | 19,505 | 18,332 (118.21) | 5,199 | 13,133 | 10,309 | 0.2836 | 0.3252 | 0.3073 |
| **[Human HG38]** | | | | | | | | | |
| Test 8 (Reference: HG38) | | | | | | | | | |
| Bayesembler | 17,669 | 15,424 | 13,919 (78.78) | 7,524 | 6,395 | 10,145 | **0.5406** | 0.4258 | 0.4764 |
| Cufflinks | 17,669 | 15,135 | 14,923 (84.46) | 7,280 | 7,643 | 10,389 | 0.4878 | 0.4120 | 0.4467 |
| Scallop | 17,669 | 29,325 | 26,857 (152.00) | 8,642 | 18,215 | 9,027 | 0.3218 | 0.4891 | 0.3882 |
| StringTie2 | 17,669 | 23,623 | 19,353 (109.53) | 8,976 | 10,377 | 8,693 | 0.4638 | **0.5080** | **0.4849** |
| Test 9 (Reference: HX1) | | | | | | | | | |
| Bayesembler | 17,669 | 15,424 | 14,610 (82.69) | 5,413 | 9,197 | 12,256 | **0.3705** | 0.3064 | 0.3354 |
| Cufflinks | 17,669 | 16,554 | 16,258 (92.01) | 5,296 | 10,962 | 12,373 | 0.3257 | 0.2997 | 0.3122 |
| Scallop | 17,669 | 19,980 | 18,779 (106.28) | 6,132 | 12,646 | 11,536 | 0.3266 | 0.3471 | **0.3365** |
| StringTie2 | 17,669 | 21550 | 20,202 (114.34) | 6,217 | 13,985 | 11,452 | 0.3077 | **0.3519** | 0.3283 |

[a]The best performance score for each dataset is shown in red boldface.
[b]Total number of transcripts in the benchmark transcriptome.
[c]Number of all contigs produced by the assembler.
[d]Number of unique contigs produced by the assembler. Proportion (%) of the number of transcripts in the benchmark transcriptome is shown in parentheses.

**Table S8. Performance analysis of genome-guided assemblers using the STAR aligner.**

| Assembler | Actual[a] | Total[b] | Unique[c] | *TP* | *FP* | *FN* | Precision | Recall | *F* |
|---|---|---|---|---|---|---|---|---|---|
| **[No0-NoAlt]** | | | | | | | | | |
| Test 4 (Reference: No-0) | | | | | | | | | |
| Bayesembler | 18,875 | - | - | - | - | - | - | - | - |
| Cufflinks | 18,875 | 21,327 | 20,925 (110.86) | 13,769 | 7,156 | 5,106 | 0.6580 | 0.7295 | 0.6919 |
| Scallop | 18,875 | 19,714 | 19,660 (104.16) | 15,652 | 4,008 | 3,223 | 0.7961 | 0.8292 | 0.8124 |
| StringTie2 | 18,875 | 19,920 | 19,668 (104.20) | 15,930 | 3,738 | 2,945 | **0.8099** | **0.8440** | **0.8266** |
| Test 5 (Reference: Col-0) | | | | | | | | | |
| Bayesembler | 18,875 | - | - | - | - | - | - | - | - |
| Cufflinks | 18,875 | 21,722 | 21,178 (112.20) | 5,953 | 15,225 | 12,922 | 0.2811 | 0.3154 | 0.2973 |
| Scallop | 18,875 | 20,385 | 20,264 (107.36) | 7,092 | 13,172 | 11,783 | 0.3500 | **0.3757** | 0.3624 |
| StringTie2 | 18,875 | 18,150 | 17,212 (91.19) | 6,625 | 10,587 | 12,250 | **0.3849** | 0.3510 | **0.3672** |
| **[Col0-Alt]** | | | | | | | | | |
| Test 6 (Reference: Col-0) | | | | | | | | | |
| Bayesembler | 15,508 | - | - | - | - | - | - | - | - |
| Cufflinks | 15,508 | 18,060 | 17,414 (112.29) | 7,959 | 9,455 | 7,549 | 0.4570 | 0.5132 | 0.4835 |
| Scallop | 15,508 | 18,188 | 17,179 (110.78) | 10,468 | 6,711 | 5,040 | **0.6093** | **0.6750** | **0.6405** |
| StringTie2 | 15,508 | 18,018 | 17,000 (109.62) | 10,061 | 6,939 | 5,447 | 0.5918 | 0.488 | 0.6190 |
| Test 7 (Reference: No-0) | | | | | | | | | |
| Bayesembler | 15,508 | - | - | - | - | - | - | - | - |
| Cufflinks | 15,508 | 18,218 | 17,441 (112.46) | 3,977 | 13,464 | 11,531 | 0.2280 | 0.2564 | 0.2414 |
| Scallop | 15,508 | 18,176 | 17,219 (111.03) | 5,293 | 11,926 | 10,215 | **0.3074** | **0.3413** | **0.3235** |
| StringTie2 | 15,508 | 18,515 | 17,477 (112.70) | 5,169 | 12,308 | 10,339 | 0.2958 | 0.3333 | 0.3134 |
| **[Human HG38]** | | | | | | | | | |
| Test 8 (Reference: HG38) | | | | | | | | | |
| Bayesembler | 17,669 | - | - | - | - | - | - | - | - |
| Cufflinks | 17,669 | 15,206 | 14,495 (82.04) | 7,628 | 6,867 | 10,041 | 0.5263 | 0.4317 | 0.4743 |
| Scallop | 17,669 | 17,940 | 17,054 (96.52) | 9,309 | 7,745 | 8,360 | **0.5459** | 0.5269 | 0.5362 |
| StringTie2 | 17,669 | 22,546 | 18,237 (103.21) | 9,761 | 8,476 | 7,908 | 0.5352 | **0.5524** | **0.5437** |
| Test 9 (Reference: HX1) | | | | | | | | | |
| Bayesembler | 17,669 | - | - | - | - | - | - | - | - |
| Cufflinks | 17,669 | 17,405 | 16,438 (93.03) | 5,491 | 10,947 | 12178 | 0.3340 | 0.3108 | 0.3220 |
| Scallop | 17,669 | 19,499 | 18,477 (104.57) | 6,557 | 11,920 | 11112 | **0.3549** | 0.3711 | 0.3628 |
| StringTie2 | 17,669 | 20,587 | 19,309 (109.28) | 6,757 | 12,552 | 10,912 | 0.3499 | **0.3824** | **0.3655** |

[a]The best performance score for each dataset is shown in red boldface.
[b]Total number of transcripts in the benchmark transcriptome.
[c]Number of all contigs produced by the assembler.
[d]Number of unique contigs produced by the assembler. Proportion (%) of the number of transcripts in the benchmark transcriptome is shown in parentheses.
[e]Bayesembler requires Tophat2 alignments, and as of this writing cannot run on alignments produced by STAR.

**Table S9. Performance analysis of the three ensemble transcriptome assemblers.[a]**

| Assembler | Actual[b] | Total[c] | Unique[d] | *TP* | *FP* | *FN* | Precision | Recall | *F* |
|---|---|---|---|---|---|---|---|---|---|
| Test 4: No0-Alt (Reference: No0) | | | | | | | | | |
| TransBorrow | 18,875 | 23,982 | 22,592 (119.69) | 15,689 | 6,903 | 3,186 | 0.6944 | 0.8312 | 0.7567 |
| ConSemble3+g | 18,875 | | 15,688 (83.12) | 14,200 | 1,488 | 4,675 | **0.9052** | **0.7523** | **0.8217** |
| ConSemble2+g | 18,875 | | 19,947 (105.68) | 15,819 | 4,128 | 3,056 | 0.7931 | **0.8381** | 0.8150 |
| ConSemble4g | 18,875 | | 9,922 (52.57) | 9,416 | 506 | 9,459 | **0.9490** | 0.4989 | 0.6540 |
| Test 6: Col0-Alt (Reference: Col0) | | | | | | | | | |
| TransBorrow | 15,508 | 1,819 | 1,800 (11.61) | 1,019 | 781 | 14,489 | 0.5661 | 0.0657 | 0.1177 |
| ConSemble3+g | 15,508 | 13,380 | 13,380 (86.28) | 9,679 | 3,701 | 5,829 | **0.7234** | **0.6241** | **0.6701** |
| ConSemble2+g | 15,508 | 17,200 | 17,200 (110.91) | 10,754 | 6,446 | 4,754 | 0.6252 | **0.6934** | 0.6576 |
| ConSemble4g | 15,508 | 7,968 | 7,968 (51.38) | 6,328 | 1,640 | 9,180 | **0.7942** | 0.4080 | 0.5391 |
| Test 8: Human HG38 (Reference: HG38) | | | | | | | | | |
| TransBorrow | 17,669 | 21,422 | 18,605 (105.30) | 8,823 | 9,782 | 8,846 | 0.4742 | **0.4993** | 0.4865 |
| ConSemble3+g | 17,669 | 11,945 | 11,945 (67.60) | 7,744 | 4,201 | 9,925 | **0.6483** | 0.4383 | **0.5230** |
| ConSemble2+g | 17,669 | 16,127 | 16,127 (91.27) | 8,864 | 7,263 | 8,805 | 0.5496 | **0.5017** | **0.5246** |
| ConSemble4g | 17,669 | 7,713 | 7,713 (43.65) | 5,670 | 2,043 | 11,999 | **0.7351** | 0.3209 | 0.4468 |

[a]The better performance score for each dataset between TransBorrow and ConSemble3+g is shown in red boldface. When ConSemble2+g or ConSemble4g showed better performance than ConSemble3+g, such scores are shown in black boldface.

[b]Total number of transcripts in the benchmark transcriptome

[c]Number of all contigs produced by the assembler.

[d]Number of unique contigs produced by the assembler. Proportion (%) of the number of transcripts in the benchmark transcriptome is shown in parentheses.

**Table S10. Performance analysis of transcriptome assembly at the nucleotide level.**[a]

| Assembler | No0-NoAlt | | | | | Col0-Alt | | | | | Human HG38 | | | | |
|---|---|---|---|---|---|---|---|---|---|---|---|---|---|---|---|
| | $F$[b] | RSEM-EVAL (x$10^{-9}$) | Nucleotide $F_1$ | Contig $F_1$ | KC | $F$[b] | RSEM-EVAL (x$10^{-9}$) | Nucleotide $F_1$ | Contig $F_1$ | KC | $F$[b] | RSEM-EVAL (x$10^{-9}$) | Nucleotide $F_1$ | Contig $F_1$ | KC |
| **[*De novo*]** | | | | | | | | | | | | | | | |
| IDBA-Tran | 0.40 | -3.91 | **0.95** | 0.26 | 0.89 | 0.33 | -3.98 | 0.82 | 0.19 | 0.82 | 0.32 | -3.50 | **0.77** | 0.14 | 0.67 |
| rnaSPAdes | 0.43 | -2.44 | 0.71 | 0.42 | 0.87 | 0.32 | -2.31 | 0.64 | 0.19 | 0.87 | 0.39 | -2.32 | 0.72 | 0.26 | 0.68 |
| SOAPdenovo-Trans | 0.46 | -2.31 | 0.92 | **0.49** | 0.86 | 0.39 | -2.28 | 0.82 | 0.36 | **0.88** | 0.30 | -2.24 | **0.77** | 0.25 | 0.68 |
| Trinity | **0.57** | **-2.30** | 0.93 | 0.48 | **0.90** | **0.53** | **-1.88** | **0.85** | **0.52** | 0.86 | **0.45** | **-1.76** | 0.75 | **0.35** | **0.70** |
| **[Genome-guided using the same reference]** | | | | | | | | | | | | | | | |
| Bayesembler | 0.66 | -3.34 | 0.78 | 0.58 | 0.71 | 0.60 | -2.86 | 0.76 | 0.59 | 0.69 | **0.48** | -2.56 | 0.72 | **0.40** | 0.72 |
| Cufflinks | **0.76** | -1.98 | **0.95** | 0.57 | 0.90 | 0.54 | -2.00 | 0.83 | 0.51 | **0.89** | 0.45 | -2.72 | 0.81 | 0.34 | 0.76 |
| Scallop | 0.75 | -1.69 | 0.88 | **0.74** | **0.92** | **0.63** | -1.62 | **0.86** | **0.67** | 0.86 | 0.39 | -1.99 | 0.73 | 0.35 | 0.85 |
| StringTie2 | 0.75 | **-0.97** | 0.90 | 0.63 | 0.91 | 0.60 | **-0.96** | 0.83 | 0.61 | 0.84 | **0.48** | **-1.05** | **0.76** | 0.38 | **0.86** |
| **[Ensemble]** | | | | | | | | | | | | | | | |
| Concatenation | 0.52 | -1.93 | 0.83 | 0.53 | 0.86 | 0.51 | -1.98 | 0.79 | **0.51** | 0.81 | 0.38 | -2.62 | 0.68 | 0.30 | 0.57 |
| EvidentialGene | 0.26 | -2.08 | 0.36 | 0.18 | 0.71 | 0.24 | -2.56 | 0.50 | 0.17 | 0.72 | 0.16 | -2.49 | 0.47 | 0.07 | 0.57 |
| TransBorrow | 0.76 | **-0.88** | **0.84** | **0.58** | **0.91** | 0.12 | -4.77 | 0.18 | 0.15 | 0.11 | 0.49 | **-1.03** | **0.73** | **0.38** | **0.85** |
| ConSemble3+d | 0.68 | -3.29 | 0.82 | 0.21 | 0.80 | 0.56 | -3.78 | 0.78 | 0.14 | 0.73 | 0.49 | -3.59 | 0.68 | 0.08 | 0.59 |
| ConSemble3+g | **0.82** | -1.88 | 0.70 | 0.51 | 0.65 | **0.67** | **-1.67** | 0.73 | 0.50 | 0.62 | **0.52** | -1.93 | 0.68 | 0.33 | 0.60 |
| ConSemble3+dLong[c] | 0.68 | -2.18 | **0.84** | 0.54 | 0.86 | 0.56 | -1.96 | 0.81 | 0.48 | **0.83** | 0.49 | -1.81 | 0.66 | 0.32 | 0.67 |
| ConSemble3+dHigh[d] | 0.68 | -2.27 | **0.84** | 0.51 | 0.85 | 0.56 | -2.05 | **0.82** | 0.45 | 0.82 | - | - | - | - | - |

[a]The best score among the assemblers for each group is shown in red boldface.
[b]From Table S3 and Tables S6 (for *de novo* assemblers) and Tests 4, 6, and 8 in Tables S7 and S9 (for genome-guided assemblers).
[c]Longest contigs producing the protein sequences kept by ConSemble are chosen.
[d]Contigs with the highest RSEM-EVAL producing the protein sequences kept by ConSemble are chosen.

**Table S11. Performance analysis for the No0-NoAlt assembly using different identity thresholds.[a]**

| Assembler | *TP* | | | Precision | | | Recall | | | *F* | | |
|---|---|---|---|---|---|---|---|---|---|---|---|---|
| | 95% | 98% | 100% | 95% | 98% | 100% | 95% | 98% | 100% | 95% | 98% | 100% |
| **[Genome-guided using the same reference]** | | | | | | | | | | | | |
| Bayesembler | 11,493 | 11,390 | 11,200 | 0.7575 | 0.7507 | 0.7382 | 0.6089 | 0.6034 | 0.5934 | 0.6751 | 0.6691 | 0.6579 |
| Cufflinks | 15,134 | 14,871 | 14,531 | **0.7846** | **0.7710** | **0.7534** | 0.8018 | 0.7879 | 0.7699 | 0.7931 | 0.7793 | **0.7615** |
| Scallop | 15,537 | 15,433 | **15,184** | 0.7261 | 0.7213 | 0.7096 | 0.8232 | 0.8176 | **0.8045** | 0.7716 | 0.7664 | 0.7541 |
| StringTie | **16,304** | **15,812** | 15,135 | 0.7530 | 0.7303 | 0.6990 | **0.8638** | **0.8377** | 0.8019 | **0.8046** | **0.7803** | 0.7469 |
| **[Genome-guided using the different reference]** | | | | | | | | | | | | |
| Bayesembler | 10,191 | 9,780 | 5,142 | 0.6562 | 0.6297 | **0.3311** | 0.5399 | 0.5181 | 0.2724 | 0.5924 | 0.5685 | 0.2989 |
| Cufflinks | 13,683 | 12,779 | 6,510 | **0.6863** | **0.6409** | 0.3265 | 0.7249 | 0.6770 | 0.3449 | 0.7051 | **0.6585** | 0.3355 |
| Scallop | 13,537 | 13,003 | **6,908** | 0.6071 | 0.5831 | 0.3098 | 0.7172 | 0.6889 | **0.3660** | 0.6576 | 0.6316 | **0.3356** |
| Stringtie | **14,632** | **13,548** | 6,857 | 0.6569 | 0.6082 | 0.3078 | **0.7752** | **0.7178** | 0.3633 | **0.7112** | **0.6585** | 0.3333 |
| **[*De novo*]** | | | | | | | | | | | | |
| IDBA-Tran | 10,368 | 9,120 | 8,344 | 0.4545 | 0.3998 | 0.3658 | 0.5493 | 0.4832 | 0.4421 | 0.4974 | 0.4375 | 0.4003 |
| rnaSPAdes | 13,505 | 12,452 | 10,034 | 0.4873 | 0.4493 | 0.3621 | **0.7155** | 0.6597 | 0.5316 | 0.5798 | 0.5345 | 0.4307 |
| SOAPdenovo | 11,931 | 11,643 | 11,118 | 0.3994 | 0.3897 | 0.3721 | 0.6321 | 0.6168 | 0.5890 | 0.4895 | 0.4777 | 0.4561 |
| Trinity | **12,950** | **12,588** | **12,057** | **0.5506** | **0.5352** | **0.5126** | 0.6861 | **0.6669** | **0.6388** | **0.6109** | **0.5939** | **0.5688** |
| **[Ensemble]** | | | | | | | | | | | | |
| Concatenation | 13,973 | 13,662 | 11,124 | 0.5915 | 0.5784 | 0.4709 | 0.7403 | 0.7238 | 0.5894 | 0.6576 | 0.6430 | 0.5235 |
| EvidentialGene | 15,216 | 14,493 | 12,519 | 0.1932 | 0.1840 | 0.1589 | 0.8061 | 0.7678 | 0.6633 | 0.3117 | 0.2969 | 0.2564 |
| ConSemble3+d | 13,695 | 13,600 | 13,352 | 0.6747 | 0.6700 | 0.6578 | 0.7256 | 0.7205 | 0.7074 | **0.6992** | **0.6944** | **0.6823** |
| ConSemble3+g | 14,630 | 14,519 | 14,223 | **0.9326** | **0.9255** | **0.9066** | 0.7751 | 0.7692 | 0.7535 | **0.8466** | **0.8401** | **0.8230** |
| TransBorrow | **17,154** | **16,539** | **15,689** | 0.7153 | 0.6896 | 0.6542 | **0.9088** | **0.8762** | **0.8312** | 0.8005 | 0.7718 | 0.7322 |

[a]The best score among the assemblers for each group is shown in red boldface. For *de novo* ensemble methods, the best *F* scores are shown in blue boldface.

**Table S12. Performance analysis for the Col0-Alt assembly using different identity thresholds.**[a]

| Assembler | *TP* | | | Precision | | | Recall | | | *F* | | |
|---|---|---|---|---|---|---|---|---|---|---|---|---|
| | 95% | 98% | 100% | 95% | 98% | 100% | 95% | 98% | 100% | 95% | 98% | 100% |
| **[Genome-guided using the same reference]** | | | | | | | | | | | | |
| Bayesembler | 10,403 | 9,906 | 9,158 | **0.6869** | **0.6541** | **0.6047** | 0.6708 | 0.6388 | 0.5905 | 0.6788 | 0.6464 | 0.5975 |
| Cufflinks | 10,312 | 9,525 | 8,560 | 0.6539 | 0.6040 | 0.5428 | 0.6649 | 0.6142 | 0.5520 | 0.6594 | 0.6091 | 0.5474 |
| Scallop | 12,281 | **11,615** | **10,534** | 0.6802 | 0.6433 | 0.5834 | 0.7919 | **0.7490** | **0.6793** | **0.7318** | **0.6921** | **0.6277** |
| StringTie | **12,284** | 11,370 | 10,034 | 0.6501 | 0.6017 | 0.5310 | **0.7921** | 0.7332 | 0.6470 | 0.7141 | 0.6610 | 0.5833 |
| **[Genome-guided using the different reference]** | | | | | | | | | | | | |
| Bayesembler | 9,083 | 8,422 | 4,810 | **0.5925** | **0.5494** | **0.3138** | 0.5857 | 0.5431 | 0.3102 | 0.5891 | 0.5462 | 0.3120 |
| Cufflinks | 8,846 | 7,811 | 4,321 | 0.5309 | 0.4688 | 0.2593 | 0.5704 | 0.5037 | 0.2786 | 0.5499 | 0.4856 | 0.2686 |
| Scallop | 10,224 | 9,366 | **5,315** | 0.5554 | 0.5088 | 0.2887 | 0.6593 | 0.6039 | **0.3427** | 0.6029 | **0.5523** | **0.3134** |
| Stringtie | **10,692** | **9,530** | 5,199 | 0.5482 | 0.4886 | 0.2665 | **0.6895** | **0.6145** | 0.3352 | **0.6107** | 0.5444 | 0.2970 |
| **[*De novo*]** | | | | | | | | | | | | |
| IDBA-Tran | 7,634 | 6,842 | 6,021 | 0.3733 | 0.3346 | 0.2945 | 0.4923 | 0.4412 | 0.3883 | 0.4246 | 0.3806 | 0.3349 |
| rnaSPAdes | 9,677 | 8,827 | 7,556 | 0.3073 | 0.2803 | 0.2399 | 0.6240 | 0.5692 | 0.4872 | 0.4118 | 0.3756 | 0.3215 |
| SOAPdenovo | 8,940 | 8,222 | 7,281 | 0.4183 | 0.3847 | 0.3407 | 0.5765 | 0.5302 | 0.4695 | 0.4848 | 0.4459 | 0.3948 |
| Trinity | **11,147** | **10,457** | **9,252** | **0.5743** | **0.5387** | **0.4767** | **0.7188** | **0.6743** | **0.5966** | **0.6385** | **0.5989** | **0.5299** |
| **[Ensemble]** | | | | | | | | | | | | |
| Concatenation | 10,463 | 9,744 | 8,487 | 0.5906 | 0.5500 | 0.4790 | 0.6747 | 0.6283 | 0.5473 | 0.6298 | 0.5865 | 0.5109 |
| EvidentialGene | 10,083 | 8,967 | 7,218 | 0.2301 | 0.2046 | 0.1647 | 0.6502 | 0.5782 | 0.4654 | 0.3399 | 0.3023 | 0.2433 |
| ConSemble3+d | 10,777 | 10,184 | 9,189 | 0.6214 | 0.5872 | 0.5299 | 0.6949 | 0.6567 | 0.5927 | **0.6561** | **0.6200** | **0.5600** |
| ConSemble3+g | **11,337** | **10,752** | **9,793** | **0.8473** | **0.8036** | **0.7319** | **0.7310** | **0.6933** | **0.6315** | **0.7849** | **0.7444** | **0.6780** |
| TransBorrow | 1,358 | 1,228 | 1,019 | 0.7466 | 0.6751 | 0.5602 | 0.0876 | 0.0792 | 0.0657 | 0.1567 | 0.1417 | 0.1176 |

[a]The best score among the assemblers for each group is shown in red boldface. For *de novo* ensemble methods, the best *F* scores are shown in blue boldface.

**Table S13. Performance analysis for the Human HG38 assembly using different identity thresholds.**[a]

| Assembler | *TP* | | | Precision | | | Recall | | | *F* | | |
|---|---|---|---|---|---|---|---|---|---|---|---|---|
| | 95% | 98% | 100% | 95% | 98% | 100% | 95% | 98% | 100% | 95% | 98% | 100% |
| **[Genome-guided using the same reference]** | | | | | | | | | | | | |
| Bayesembler | 8,881 | 8,533 | 7,524 | **0.6380** | **0.6130** | **0.5405** | 0.5026 | 0.4829 | 0.4258 | **0.5623** | **0.5403** | **0.4764** |
| Cufflinks | 8,690 | 8,209 | 7,280 | 0.5823 | 0.5501 | 0.4878 | 0.4918 | 0.4646 | 0.4120 | 0.5333 | 0.5037 | 0.4467 |
| Scallop | 10,175 | 9,761 | 8,642 | 0.3788 | 0.3634 | 0.3218 | 0.5759 | 0.5524 | 0.4891 | 0.4570 | 0.4384 | 0.3882 |
| StringTie | **11,166** | **10,460** | **8,976** | 0.4727 | 0.4428 | 0.3800 | **0.6320** | **0.5920** | **0.5080** | 0.5408 | 0.5066 | 0.4348 |
| **[Genome-guided using the different reference]** | | | | | | | | | | | | |
| Bayesembler | 7,486 | 7,061 | 5,413 | **0.5124** | **0.4833** | **0.3705** | 0.4237 | 0.3996 | 0.3064 | **0.4638** | **0.4375** | 0.3354 |
| Cufflinks | 7,399 | 6,855 | 5,296 | 0.4551 | 0.4216 | 0.3257 | 0.4188 | 0.3880 | 0.2997 | 0.4362 | 0.4041 | 0.3122 |
| Scallop | 8,406 | 7,938 | 6,132 | 0.4476 | 0.4227 | 0.3265 | 0.4757 | 0.4493 | 0.3470 | 0.4613 | 0.4356 | **0.3365** |
| Stringtie | **8,888** | **8,199** | **6,217** | 0.4124 | 0.3805 | 0.2885 | **0.5030** | **0.4640** | **0.3519** | 0.4532 | 0.4181 | 0.3170 |
| **[*De novo*]** | | | | | | | | | | | | |
| IDBA-Tran | 7,750 | 7,177 | 6,154 | 0.3698 | 0.3425 | 0.2937 | 0.4386 | 0.4062 | 0.3483 | 0.4013 | 0.3716 | 0.3187 |
| rnaSPAdes | 9,727 | 9,034 | 7,637 | 0.4579 | 0.4252 | 0.3595 | 0.5505 | 0.5113 | 0.4322 | 0.4999 | 0.4643 | 0.3925 |
| SOAPdenovo | 7,614 | 7,071 | 5,933 | 0.3460 | 0.3213 | 0.2696 | 0.4309 | 0.4002 | 0.3358 | 0.3838 | 0.3564 | 0.2991 |
| Trinity | **10,385** | **9,980** | **8,764** | **0.4881** | **0.4690** | **0.4119** | **0.5878** | **0.5648** | **0.4960** | **0.5333** | **0.5125** | **0.4500** |
| **[Ensemble]** | | | | | | | | | | | | |
| Concatenation | 7,497 | 7,078 | 6,032 | 0.5332 | 0.5034 | 0.4290 | 0.4243 | 0.4006 | 0.3414 | 0.4725 | 0.4461 | 0.3802 |
| EvidentialGene | 8,775 | 7,895 | 6,248 | 0.1482 | 0.1333 | 0.1055 | 0.4966 | 0.4468 | 0.3536 | 0.2282 | 0.2054 | 0.1625 |
| ConSemble3+d | 10,673 | **10,315** | **9,128** | 0.5516 | 0.5331 | 0.4717 | 0.6041 | **0.5838** | **0.5166** | **0.5766** | **0.5573** | **0.4932** |
| ConSemble3+g | 9,447 | 9,066 | 8,045 | **0.7909** | **0.7590** | **0.6735** | 0.5347 | 0.5131 | 0.4553 | **0.6380** | **0.6123** | **0.5433** |
| TransBorrow | **10,982** | 10,273 | 8,823 | 0.5127 | 0.4796 | 0.4119 | **0.6215** | 0.5814 | 0.4993 | 0.5619 | 0.5256 | 0.4514 |

[a]The best score among the assemblers for each group is shown in red boldface. For *de novo* ensemble methods, the best *F* scores are shown in blue boldface.
