## Additional file 3 for "A consensus-based ensemble approach to improve transcriptome assembly"

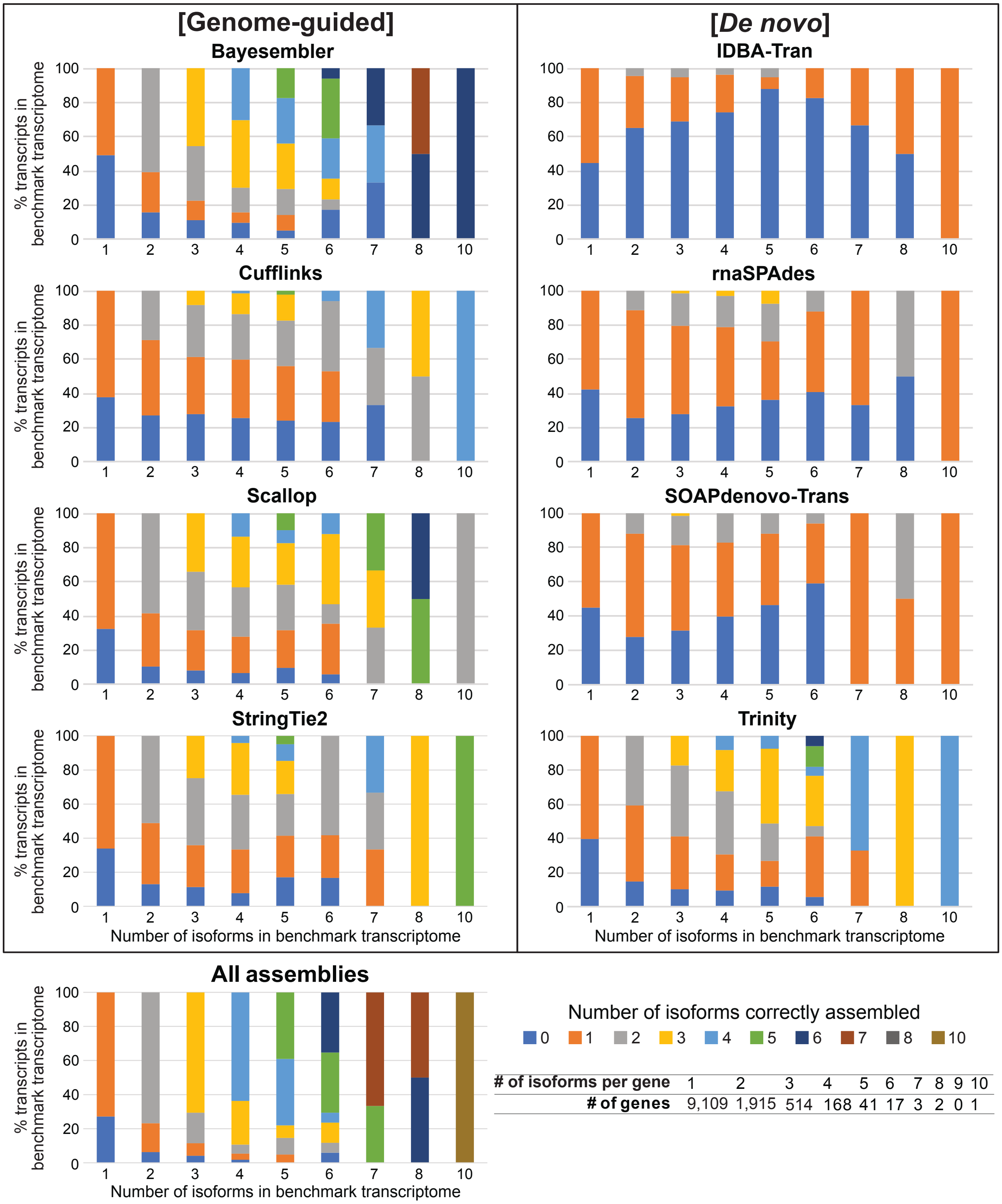


**Fig. S1. Proportion (%) of correctly assembled isoforms per gene by different assemblers.** Genes are grouped by the number of the isoforms existing in the reference. For each group, the proportion for each assembled isoform number (from 0 to 9) is color coded where 0 indicates no transcripts were assembled for a given gene. Results are based on Test 3 (Col0-Alt using the Col-0 reference) for genome-guided and Test 8 (Col0-Alt) for *de novo* assembly (with default settings) in Table S2 in Additional file 2. In "All assemblies", all genome-guided and *de novo* assemblies across all kmer lengths were merged.

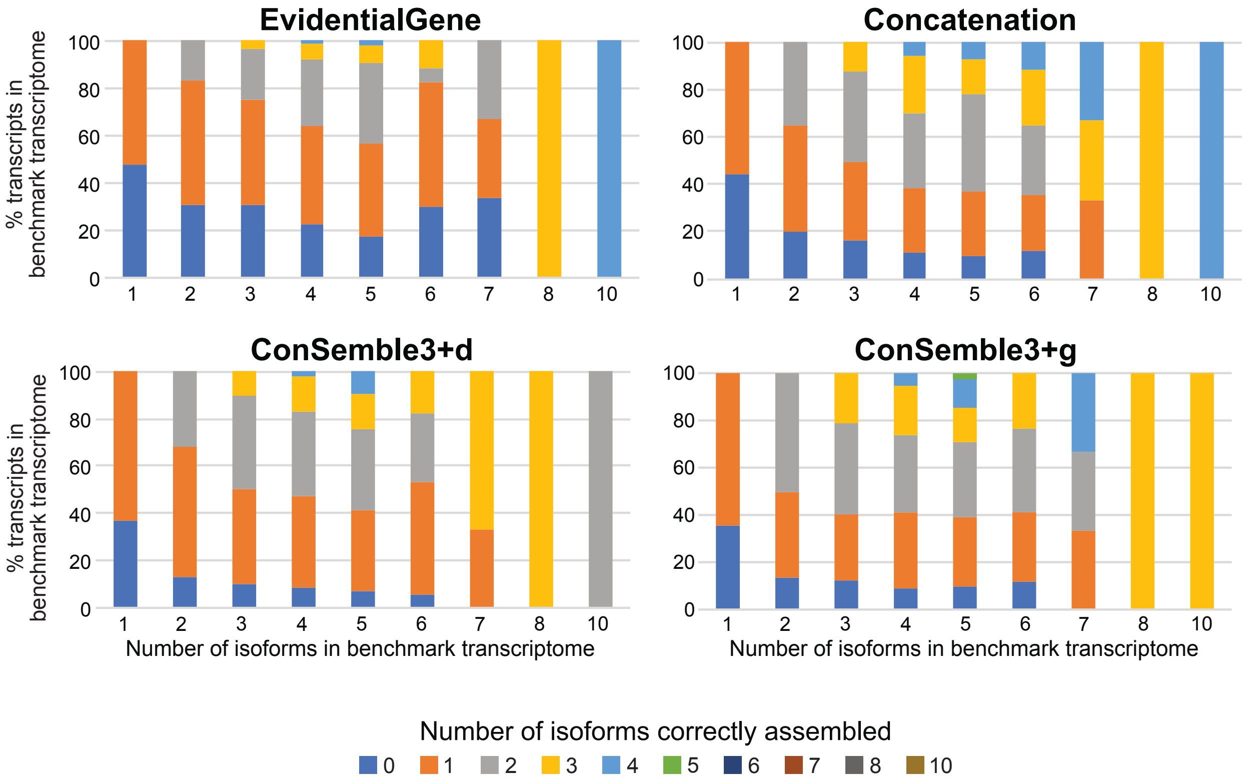


| # of isoforms per gene | 1 | 2 | 3 | 4 | 5 | 6 | 7 | 8 | 9 | 10 |
|---|---|---|---|---|---|---|---|---|---|---|
| # of genes | 9,109 | 1,915 | 514 | 168 | 41 | 17 | 3 | 2 | 0 | 1 |

**Fig. S2. Proportion of correctly assembled isoforms per gene by different ensemble assemblers.** Genes are grouped by the number of isoforms existing in the reference. For each group, the proportion for each assembled isoform number (from 0 to 9) is color coded where 0 indicates no transcripts were assembled for a given gene. Results are based on Test 3 (Col0-Alt using Col-0 reference) for ConSemble3+g and Test 8 (Col0-Alt) for EvidentialGene, Concatenation, and ConSemble3+d (see Table S2 in Additional file 2).

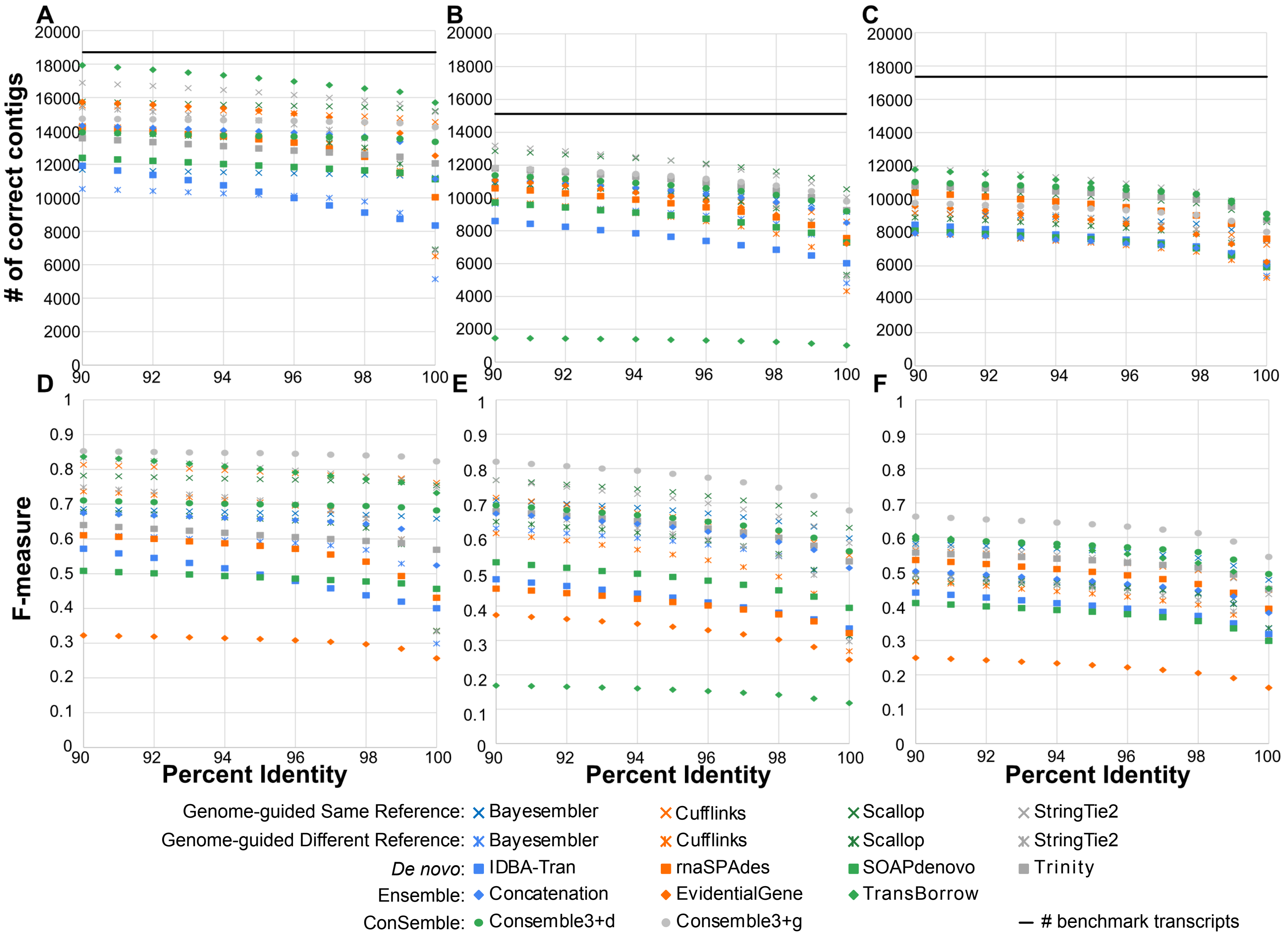


**Fig. S3. The numbers of correctly assembled contigs with varied percent identity threshold for true positives.** The threshold % identity was varied from 90% to 100% (at the protein level). The number of correctly assembled contigs was counted for each % identity threshold (A, B, and C). These correctly assembled contigs for each threshold were also used to calculate the F-measure (D, E, and F). The dataset used was No0-NoAlt (A and D), Col0-Alt (B and E), or Human HG38 (C and F). See Tables S11-S13 in Additional file 2 for more statistics.

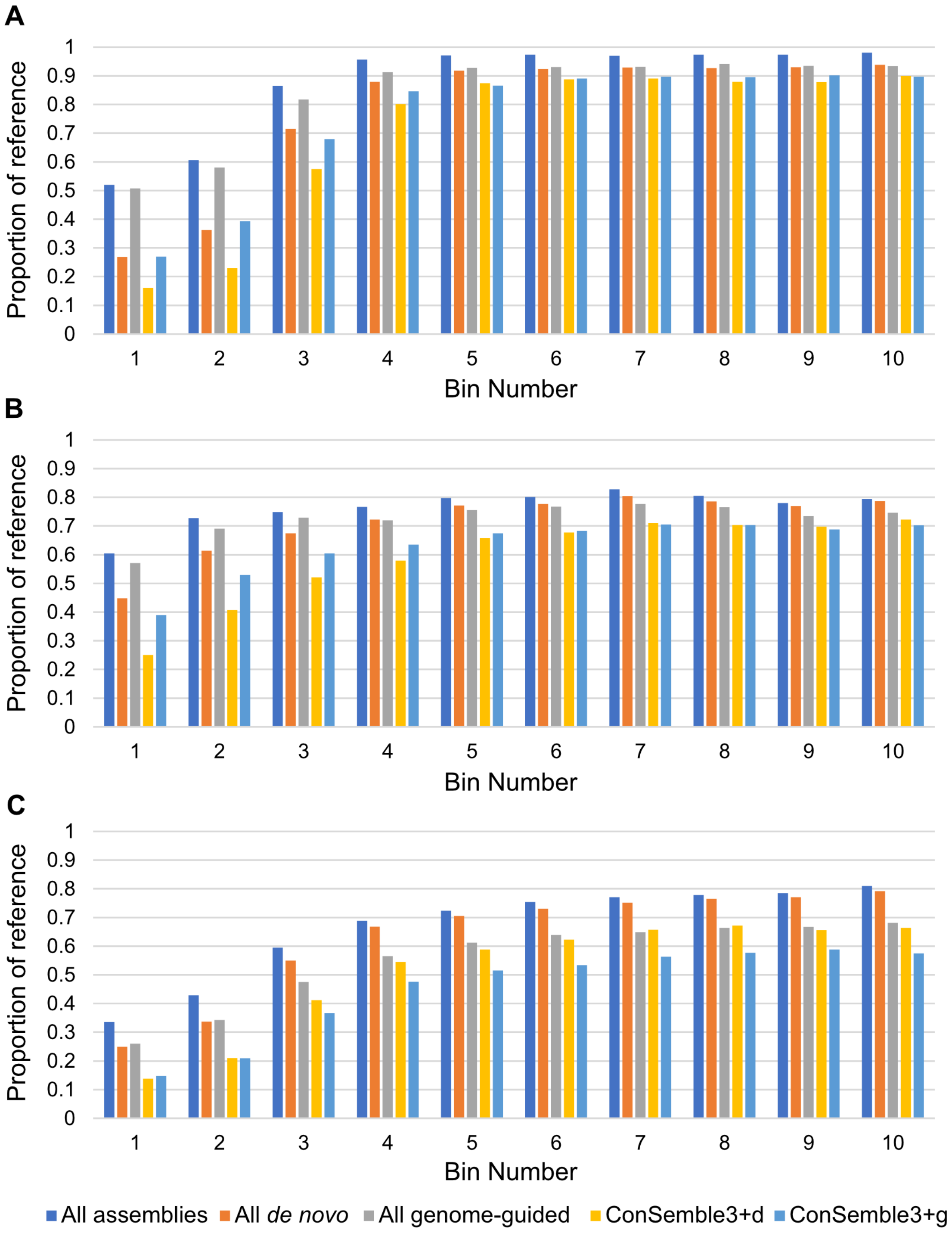


**Fig. S4. Proportion of correctly assembled transcripts by expression.** Genes are grouped into ten equally sized bins by expression decile in the benchmark datasets (A: No0-NoAlt, B: Col0-Alt, and C: Human). For each bin, the proportion of benchmark transcripts correctly assembled for each of the following categories are shown: "All assemblies": transcript is correctly assembled by any *de novo* or genome-guided methods; "All *de novo*": transcript is correctly assembled by any *de novo* method, regardless of the kmer length; "All genome-guided": transcript is correctly assembled by any genome-guided method; "ConSemble3+d": transcript is correctly assembled by ConSemble3+d; and "ConSemble3+g": transcript is correctly assembled by ConSemble3+g.
